# Comprehensive identification of β-lactam antibiotic polypharmacology in *Mycobacterium tuberculosis*

**DOI:** 10.1101/2025.02.25.640225

**Authors:** Kaylyn L. Devlin, Emily Hutchinson, Hailey N. Dearing, Samantha R. Levine, Deseree J. Reid, Damon T. Leach, Lydia H. Griggs, Gerard X. Lomas, Leo J. Gorham, Aaron T. Wright, Gyanu Lamichhane, Vivian S. Lin, Kimberly E. Beatty

## Abstract

Infections with *Mycobacterium tuberculosis* (*Mtb*) cause tuberculosis (TB), which requires at least six months of treatment with multiple antibiotics. There is emergent interest in using β-lactam antibiotics to improve treatment outcomes for patients. These drugs target cell wall biosynthesis, but a comprehensive list of enzymes inhibited by β-lactams in *Mtb* is lacking. In the current study, we sought to identify and characterize *Mtb* enzymes inhibited by β-lactam antibiotics using physiological conditions representative of both acute and chronic TB disease. We used new activity-based probes based on the β-lactam antibiotic meropenem due to its approval by the World Health Organization for TB treatment. Activity-based probes label enzymes based on both substrate specificity and catalytic mechanism, enabling precise identification of drug targets. We identified previously undiscovered targets of meropenem in addition to known cell wall biosynthetic enzymes. We validated β-lactam binding and hydrolysis for six newly identified targets: Rv1723, Rv2257c, Rv0309, DapE (Rv1202), MurI (Rv1338), and LipD (Rv1923). Our results demonstrate that there are at least 30 enzymes in *Mtb* vulnerable to inhibition by meropenem. This is many more β-lactam targets than historically described, suggesting that efficacy in *Mtb* is a direct result of polypharmacology.

## Introduction

Tuberculosis (TB) is the most deadly infectious disease in human history with annual infection rates of over 10 million people [1]. TB is caused by *Mycobacterium tuberculosis* (*Mtb*), a pathogen that is highly adept at surviving in the human host. *Mtb* can respond to adverse conditions by switching to a dormant, non-replicating state, enabling it to persist in the human body for years as a latent infection. Even people with active TB host multiple sub-populations of *Mtb* across a spectrum from actively replicating to dormant[2-4]. The recommended treatment for TB is six months of therapy with four drugs (i.e., isoniazid, rifampicin, pyrazinamide, and ethambutol). The long duration of treatment is in part because it is difficult to eradicate dormant *Mtb* “persisters”[5], which have phenotypic resistance to drugs[6, 7]. Additionally, some *Mtb* strains have developed resistance to some or all anti-mycobacterial drugs[8-10], making efficacious treatment even more challenging. New safe and accessible treatment regimens that can address these challenges are urgently needed[1, 11].

We and others have studied the potential of repurposing clinically-approved β-lactam antibiotics to treat TB and drug-resistant TB[12-20]. Currently, no first-line TB drug is a β-lactam antibiotic, and outdated assumptions have hampered broad consideration of their use. One assumption is that *Mtb*’s expression of the extended-spectrum Class A β-lactamase BlaC (Rv2068c) renders β-lactams ineffective. Yet BlaC has limited activity against carbapenems[21-26] and is inactivated by the clinically-approved β-lactamase inhibitor clavulanate[27]. This means that BlaC activity can be neutralized through treatment with certain drugs.

In fact, there is clinical and experimental evidence that TB can be successfully treated with a β-lactam plus clavulanate [21, 28-38]. A 1999 report described using amoxicillin/clavulanate to cure multi-drug resistant TB in patients[36]. More recently, carbapenems (e.g., imipenem[34] and meropenem/clavulanate[29-31, 35]) were used to treat drug-resistant TB. Meropenem/clavulanate was also effective against non-replicating *Mtb in vitro*[21, 23]. Some drug-resistant strains of *Mtb* have enhanced susceptibility to β-lactam antibiotics[37, 38], making these drugs appealing for treating drug-resistant TB. For several years, the World Health Organization (WHO) has conditionally recommended the use of meropenem/clavulanate in treatment regimens for patients with drug-resistant TB due to meropenem’s superior potency among β-lactams[39]. One outcome of these studies is an interest in identifying how β-lactams can be leveraged to improve TB treatment.

A generalized model describing the mechanism of action for β-lactam antibiotics originated from evidence collected in well-studied bacteria, such as *Escherichia coli* and *Bacillus subtilis*. In these bacteria, the cell wall peptidoglycan (PG) is cross-linked by D,D-transpeptidases (DDTs), also called penicillin-binding proteins (PBPs)[40]. β-lactams were found to inhibit PBPs by mimicking the D-Ala-D-Ala portion of the PG peptide stem[41], thus targeting bacterial cell wall biosynthesis[12, 14]. Based on these early studies in model organisms, it was assumed for decades that the targets of β-lactams in mycobacteria were solely PBPs. Indeed, there is experimental evidence that β-lactams inhibit *Mtb*’s PBPs, including carboxypeptidases (DacB[42, 43], DacB1[42, 44], DacB2[42, 44, 45]] and DDTs [PonA1[46, 47], PonA2[48, 49], PbpA[50], PbpB[51]).

However, the assumption that β-lactam efficacy against *Mtb* is exclusively due to inhibition of PBPs was incomplete. In *Mtb* and other mycobacteria, the PG cross-links are formed primarily by L,D-transpeptidases (**LDT**s), with a minority of linkages formed by DDTs[52-55]. This suggests that in mycobacteria the LDTs are at least as important a drug target as the PBPs. There are five known mycobacterial LDTs (Ldt1[53, 56], Ldt2[56-58], Ldt3[56], Ldt4[56], and Ldt5[59]). They are evolutionarily distinct from PBPs and share no sequence or structure homology. The PBPs have a catalytic serine within a characteristic SXXK motif while the LDTs have a catalytic cysteine within a conserved YkuD domain[53, 56]. An effective drug targeting mycobacterial PG biosynthesis would need to inhibit both PBPs and LDTs at once, exhibiting polypharmacology. The carbapenem class of β-lactams is unique in this regard, as they strongly inhibit both enzyme classes[56, 58-62].

Given this polypharmacology, it is possible that β-lactam efficacy against *Mtb* is dependent on the inhibition of additional, unidentified targets. To our knowledge, prior studies have collectively identified 22 *Mtb* enzymes that interact with β-lactams to varying extents. This includes the PBPs and LDTs and the β-lactamase BlaC. Other *Mtb* enzymes that hydrolyze β-lactams include Pbp-lipo[42], Rv0406[63], Rv3677c[63], Rv1367c[42], Rv1730c[42], RNase J [64], LipE [65], LipL [66], and CrfA[67]. Overall, our understanding of the mechanistic basis for β-lactam activity against *Mtb* remains incomplete. This knowledge is essential for fully harnessing the potential of β-lactams to effectively treat TB. Drug selection and optimization would benefit from a comprehensive list of targets that are active in physiologically-relevant host-pathogen conditions.

Activity-based probes (ABPs) are ideal tools for evaluating drug-enzyme interactions, as well as enzyme function and activity[68]. These small molecule probes can be designed around an inhibitor backbone to covalently label target enzymes based on catalytic mechanism. ABPs have enabled new discoveries in a variety of pathogens, including *B. subtilis*[69], *Staphylococcus aureus*[70, 71], *Enterococcus faecium*[72], and *Mtb*[17, 73-82]. For example, we previously used a set of fluorescent ABPs to image β-lactam targets, including PBPs and LDTs, in protein gel-resolved lysates of *Mtb*[17]. In that work, fluorescent β-lactams were used to discover that β-lactam targets underwent substantial changes during hypoxia-induced dormancy. A limitation of that work was that only some β-lactam targets were identified.

The current work describes the comprehensive identification of β-lactam targets in *Mtb* using a method termed activity-based protein profiling (ABPP). We used a new biotinylated ABP, meropenem-biotin (Mero-biotin), to label enzyme targets across distinct growth conditions that model environmental conditions present during active and chronic TB infection. Our results reveal previously unknown targets of β-lactams and suggest a highly promiscuous mechanism of action for carbapenems in *Mtb*. We validated a subset of novel targets and provide evidence that meropenem binds all of them. Overall, our study challenges the historical view that β-lactam activity in *Mtb* is due solely to the inhibition of PBPs.

## Results and Discussion

### Description of activity-based probes

The commercially-available fluorescent penam Bocillin-FL[83] has limited applicability for mycobacterial research for two reasons. First, penams are not recommended for use in TB patients without a companion β-lactamase inhibitor[39] because they are rapidly inactivated by BlaC hydrolysis[27]. Second, penams bind to a limited number of relevant *Mtb* β-lactam targets because they do not bind LDTs[17]. As a result, we instead selected a carbapenem, meropenem, for development of a novel ABP. Importantly, meropenem is included in the WHO recommendations for TB treatment.

Two new ABPs were designed and synthesized for the current work: meropenem-biotin (Mero-biotin) and meropenem-sulfoCy5 (Mero-sCy5) (**Figure 1A**). They were synthesized using amide bond forming conditions to attach the reporter at the secondary amine of the pyrrolidine ring of meropenem, as described[17, 69]. We made Mero-biotin for ABPP (e.g., affinity enrichment) and Mero-sCy5 for fluorescent detection of proteins by SDS-PAGE (see **Figure 1B**). We additionally used the chromogenic cephalosporin nitrocefin[84], to detect β-lactamase activity, and our previously reported Mero-Cy5[17]. Structures of these ABPs are provided in the electronic supporting information (ESI; **Scheme S1**).

**Figure 1.**
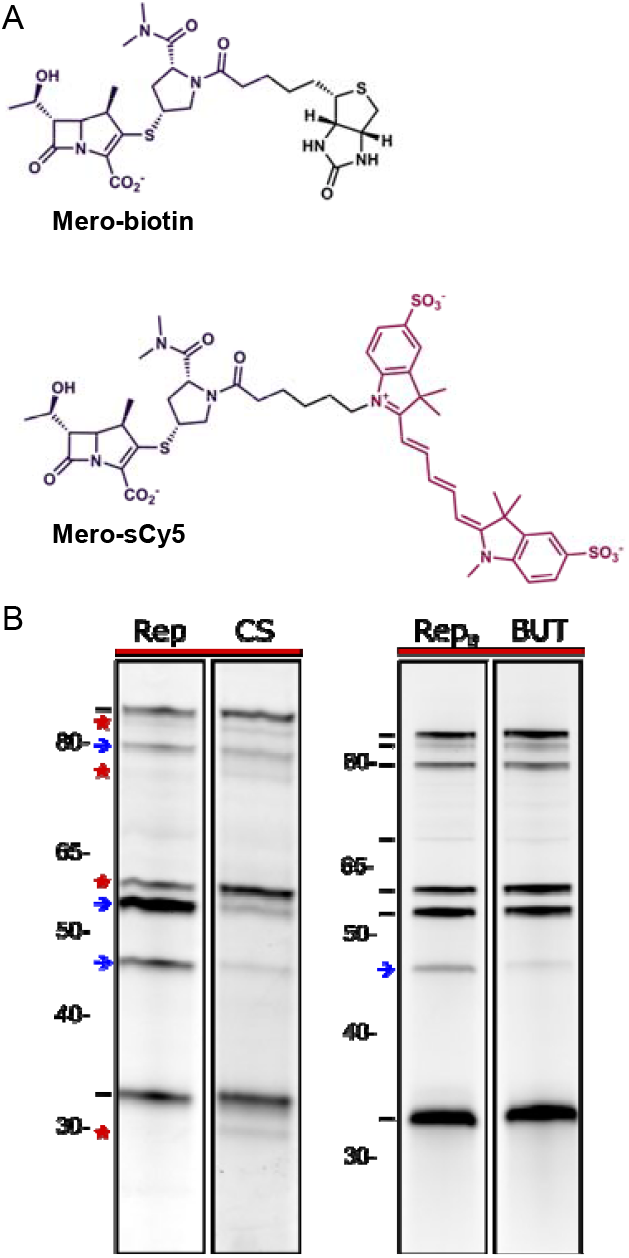
Novel ABPs enable labeling and detection of proteins that interact with β-lactams in *Mtb*. **A**. Structures of new meropenem-based ABPs, Mero-biotin and Mero-sCy5, used in the current work. **B**. Analysis of β-lactam targets in gel resolved lysates of *Mtb*. Lysates were prepared from the indicated growth conditions, treated with fluorescent meropenem, resolved by SDS-PAGE, and imaged. Legend: Black dashes indicate equivalent enzyme labeling, or Cy5 signal, between the altered growth condition and matched control. Red stars indicate increased labeling and blue arrows indicate decreased labeling in altered growth condition relative to matched control.

### Modeling host environments

There are many *in vitro* models that mimic specific host environments during acute and chronic phases of TB[85, 86]. We induced a non-replicating state using a carbon-starvation (CS) model that is known to mimic dormant infection[87, 88]. We compared the non-replicating state to active replication of *Mtb* grown under standard growth conditions (Rep). Additionally, we used a lipid-rich model (BUT) to mimic the environment found in host lung granulomas[89, 90]. This model uses the short-chain fatty acid butyrate as a carbon source in MOPS-buffered (pH 7) medium[91]. The control condition matched to BUT was grown to mid-log phase in standard medium buffered with MOPS (Rep_B_). Biological replicates (n = 6) were prepared for each condition. Inclusion of distinct conditions in our analysis enabled robust assessment of proteomic hits, as described below.

Our fluorescent ABP, Mero-sCy5, was used for a preliminary assessment of meropenem targets in *Mtb* grown under the four conditions. Lysates were resolved by SDS-PAGE and fluorescent imaging showed numerous distinct bands between 30 and 100 kDa (**Figure 1B**). Band intensities varied, indicating that some proteins differed in abundance or activity between conditions. Distinct variation was observed between Rep and CS, with some bands diminished and others enhanced in CS. Labeling patterns were similar between BUT and Rep_B_.

Identification of meropenem targets through protein gel analysis has significant limitations. First, sensitivity of target detection is relatively low, which can lead to an underestimation of the number of targets present. Second, acquiring the identity of a specific labeled target is challenging due to co-migration of many proteins in the gel. We previously encountered both issues when we used band-excision mass spectrometry (MS) analysis to identify *Mtb* proteins corresponding to fluorescent bands[17]. For example, the band appearing at ~40-45 kDa contained peptides from 14 *Mtb* proteins predicted to interact with β-lactams, including two LDTs, four PBPs, and eight putative β-lactamases. These challenges illustrate the benefits of ABPP, an unbiased approach for definitive and sensitive target identification.

### Global proteomic analysis

Alongside ABPP, we performed global proteomics analysis via liquid chromatography-tandem MS (LC-MS/MS) to determine overall abundances of proteins in each sample type. We recently described the global results from Rep and CS conditions[92]. These data allowed us to validate our enrichment protocol and determine the relative abundance of proteins before enrichment.

To summarize, we achieved 52.2% proteome coverage in at least three replicates of either CS or Rep groups. Most proteins (71.8%) were identified in both conditions, although 37% were altered in abundance in CS (≥ 2-fold change, p ≤ 0.05). Many known targets of β-lactam antibiotics were not detected in the global analysis, including most LDTs. A known challenge of global proteomics is that low abundance proteins may not be consistently observed without fractionation or other time- and cost-intensive strategies to achieve deeper coverage; ABP-based enrichment offers a solution to this problem.

### Mero-biotin activity-based protein profiling

We used ABPP to comprehensively identify meropenem targets in *Mtb* across biologically relevant states. Whole cell lysates from the four culture conditions were labeled with Mero-biotin (or mock-treated) before affinity enrichment and analysis. Targets of Mero-biotin identified by proteomics were stringently defined through the application of several criteria. First, target proteins had to be identified in at least three out of six group replicates. Second, the protein’s mean intensity value had to be significantly higher (≥ 4-fold, p ≤ 0.05) relative to mock-treated (i.e., non-probed) control samples. Most identified targets were only present in ABP-treated samples (absent in non-probed controls). Some, however, were found in non-probed controls due to specific or non-specific interaction with the streptavidin resin used for affinity enrichment, despite extensive wash steps. False hits from endogenously biotinylated proteins were expected because they are particularly abundant in *Mtb* (see ESI, **Table S1**). Any biotinylated proteins that passed the hit selection criteria were manually removed from the final target lists.

The final number of Mero-biotin targets identified in the four groups analyzed were as follows: Rep (193), CS (225), BUT (36), Rep_B_ (50). A comprehensive list of identified proteins is included in **Table S2**, and data were deposited in the Mass Spectrometry Interactive Virtual Environment (MassIVE) repository (MSV000097053). The identified targets were more numerous than anticipated, indicating that meropenem binds to many proteins beyond the expected PBPs and LDTs in whole-cell lysates. We acknowledge that it is unlikely that meropenem targets all of these proteins *in vivo*. Some may be irrelevant during infection if their β-lactam affinity is low or their localization precludes drug binding.

A direct comparison of global proteomics versus ABPP underscored several advantages of our approach (**Figure 2**). Mero-biotin enabled the enrichment and analysis of low-abundance proteins that are commonly missed in global proteomics. Enrichment of proteins based on enzymatic activity is demonstrated by the fact that patterns of protein abundance in ABPP groups do not mirror global abundance. Moreover, ABPP provides new insights into protein function and has the powerful ability to uncover novel drug targets.

**Figure 2.**
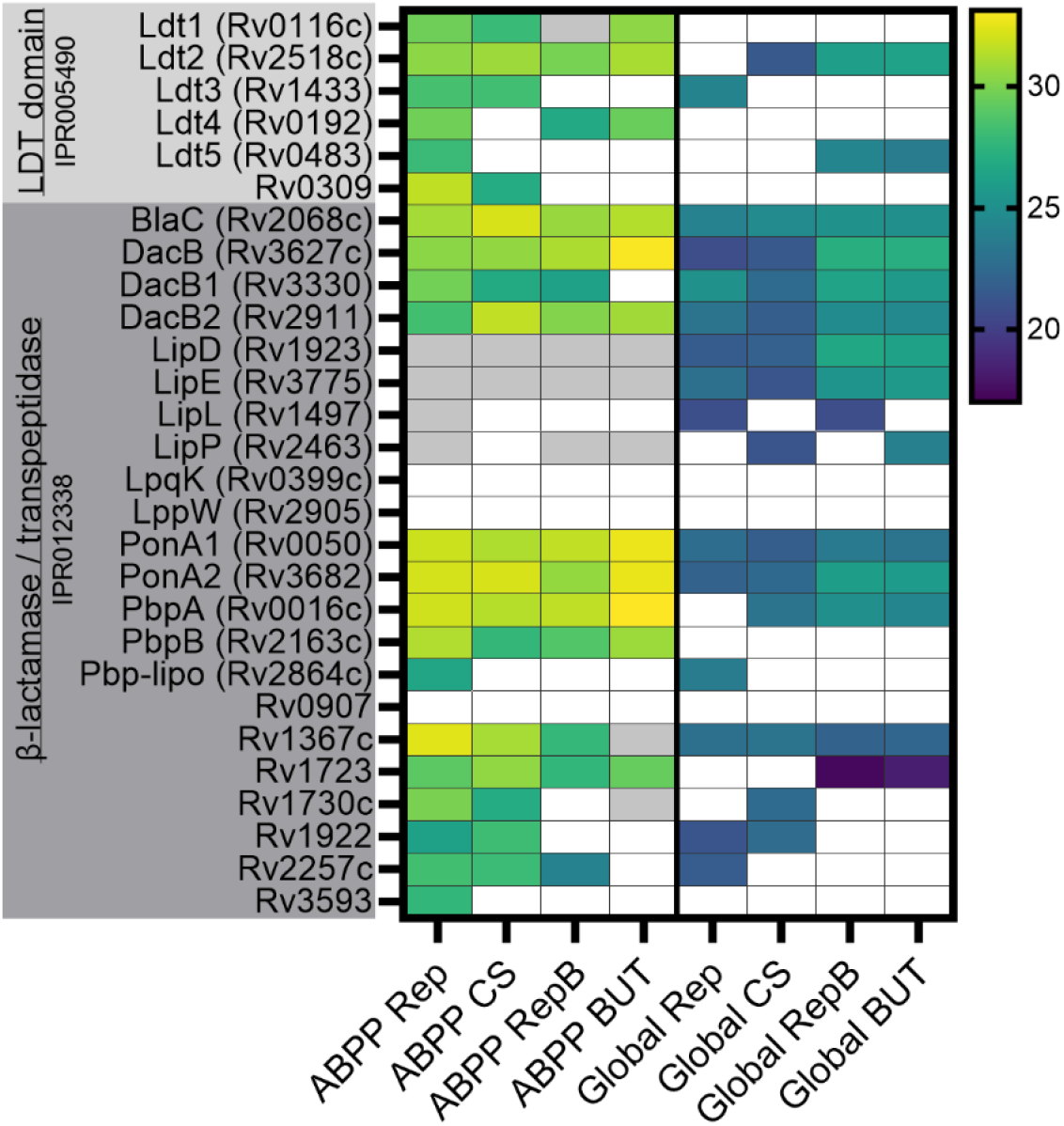
ABPP confirmation of Interpro-predicted protein functional class. Heat map of *Mtb* enzymes with Interpro protein family classification of LDT catalytic domain (IPR005490) or β-lactamase/transpeptidase-like (IPR012338). Heat map displays group mean log2 label-free quantitation intensity. A white box indicates a protein was found in n < 3 samples. A grey box indicates the protein was found in that group (n ≥ 3) but not identified as a Mero-biotin hit by our criteria.

### ABPP confirmation of predicted β-lactam-interacting proteins

We began our ABPP analysis by searching for proteins bioinformatically predicted to interact with β-lactams. We used the Interpro database[93] to identify *Mtb* enzymes predicted to have a metallo-β-lactamase (MBL) domain (IPR001279), LDT catalytic domain (IPR005490), or β-lactamase/transpeptidase-like (IPR012338) domain. *Mtb* encodes 16 proteins with putative MBL domains, including three that were verified to hydrolyze β-lactams (RNaseJ[64], Rv0406c[63], and Rv3677c[63]). None of the putative MBLs were identified in the current work. The catalytic mechanism of MBLs is metal-dependent and non-covalent. As a result, MBL interactions with Mero-biotin were unlikely to survive our enrichment protocol, which involved extensive wash steps to remove non-specifically and non-covalently bound proteins.

The other two categories, LDT catalytic domain and the β-lactamase/transpeptidase-like family, included 28 putative targets of β-lactams (**Figure 2**). Any of these proteins that were identified as meropenem targets by ABPP were considered top hits. For example, there were six proteins predicted to contain an LDT-domain, and ABPP identified all of them. Notably, we identified a previously unrecognized LDT-domain-containing enzyme, Rv0309, in both Rep and CS samples. Rv0309 has the conserved LDT active site cysteine and shares 15-20% amino acid sequence identity with the other *Mtb* LDTs. We selected Rv0309 for further characterization, as described below. Three ABPP-identified LDTs (Ldt1, Ldt4, Rv0309) were undetected in global proteomics data, highlighting the improved sensitivity obtained from ABP-enriched samples.

There were 22 β-lactamase/transpeptidase-like family members categorized in Interpro, of which we identified 15 (68%) as Mero-biotin hits (**Figure 2**). We identified four previously unknown targets of β-lactam antibiotics, namely Rv1723, Rv1922, Rv2257c, and Rv3593. The amino acid sequences of these proteins all contain a conserved Class A β-lactamase (SXXK) catalytic motif (**Figure S1**). We selected two of these putative β-lactamases for further characterization: Rv1723 and Rv2257c.

Among the enzymes in **Figure 2**, several were differentially identified in Rep versus CS. Four enzymes were missing from our CS samples, suggesting that they were down-regulated or less active in CS. The CS-absent enzymes were Ldt4 (Rv0192), Ldt5 (Rv0483), PBP-lipo (Rv2864c), and a putative β-lactamase (Rv3593).

### Mero-biotin target analysis

Presence of a β-lactam target in *Mtb* cultured across the conditions included in our study is indicative of its functional importance in both chronic and active TB disease. There were nine enzymes identified by ABPP across all conditions: PonA1, PonA2, PbpA, PbpB, DacB, DacB2, Ldt2, BlaC, and uncharacterized Rv1723. Consistent with functional importance, both DacB and PbpB are essential[94, 95], PonA2 confers a growth advantage[95], and Ldt2 is required for cell morphology, normal growth, and virulence[60].

We next identified proteins that were found as meropenem targets in at least three of four groups, and considered these top ABPP hits (**Figure 3, Table S3**). This highlighted eight additional proteins (Ldt1, Ldt4, DacB1, Rv1367, Rv2257c, DapE, Ddn, and Rv3607c), for a total of 17 top ABPP hits present in a majority of growth conditions. Of these, 12 (70%) were known PBPs, LDTs, and β-lactamases, validating the rigor of our analysis. The remaining five proteins were previously unknown as β-lactam targets, and included Rv1723, Rv2257c, Rv3706c, Ddn (Rv3547), and DapE (Rv1202). Two of these, Rv1723 and Rv2257c, were bioinformatically annotated as putative β-lactamases and were selected for further characterization. Rv3706c is a conserved hypothetical protein. Sequence analysis of Rv3706c suggests a putative *N-* terminal intracellular domain (residues 1-25) and a transmembrane domain (residues 26-48) anchoring an extracellular *C-*terminus. The final two hits were enzymes with established functions: Ddn and DapE. Ddn reduces quinones to dihydroquinones[96] and is linked to pretomanid resistance[97]. The other hit, DapE, is an essential enzyme of the succinylase pathway for the biosynthesis of both L-Lysine and *meso*-diaminopimelic acid (*meso*-DAP). The latter is integral to the *Mtb* cell wall as the central residue of the PG pentapeptide stem[98, 99]. Furthermore, orthologous DapE enzymes are metallo-hydrolases that cleave amide bonds[100], a reaction also catalyzed by β-lactamases. For these reasons, we selected DapE for further characterization.

**Figure 3.**
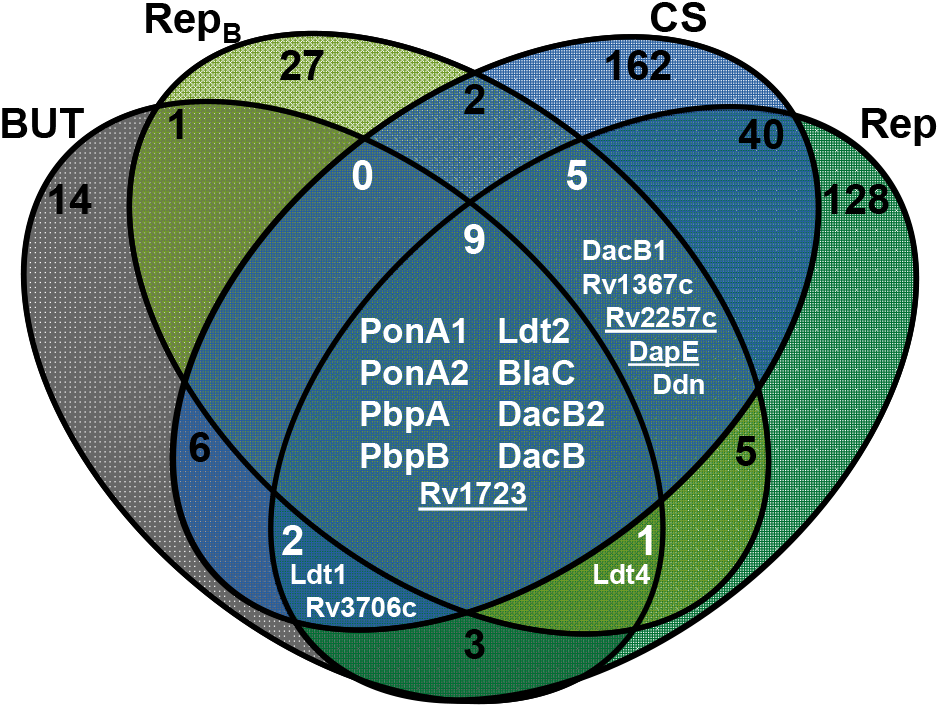
Venn diagram of *Mtb* Mero-biotin target proteins identified by ABPP in physiologically relevant growth conditions. Proteins highlighted in white were identified in at least three conditions and are considered high confidence hits. Underlined proteins were selected for further characterization.

### Selection of MurI and LipD for validation

We manually assessed the Mero-biotin targets identified by ABPP using online databases (Uniprot[101], Interpro[93], Mycobrowser[102]) and protein sequence analysis tools (AlphaFold[103], Phyre[104], and Clustal Omega[105]). As a result, we selected the glutamate racemase MurI (Rv1338) and the lipase LipD (Rv1923) for validation. MurI was identified as a Mero-biotin target in carbon starvation and lipid-rich conditions. Like the PBPs, MurI plays a key role in PG biosynthesis, converting L-Glu to D-Glu for placement in the peptide stem[106]. Glutamate racemase activity is absent in humans, but essential for *Mtb*, making MurI an attractive drug target. MurI is proposed to be localized to the membrane, suggesting it would be accessible to meropenem or other drugs. Interaction of β-lactams with MurI would be unprecedented, but MurI has nucleophilic active site residues (e.g., Cys75, Cys185, Ser13) that could potentially be acylated by a carbapenem.

LipD did not meet our ABPP hit selection criteria because it was only 2-fold enriched (p = 0.0004) in Rep samples compared to non-probed controls. However, we selected LipD because it has the β-lactamase (SXXK) catalytic motif (**Figure S1**). Two closely related *Mtb* proteins, LipE (Rv3775) and LipL (Rv1497), have β-lactamase activity[65, 66]. Lastly, we choose LipD because we have established expertise in the *Mtb* lipases[75, 82, 107].

### Validation of β-lactam binding and hydrolysis by ABPP-identified enzymes

From among the described hits, we selected six enzymes for validation: Rv1723, Rv2257c, DapE, Rv0309, MurI, and LipD. For comparison, we evaluated three additional enzymes (BlaC, DacB, Rv1367c) with established β-lactamase activity. BlaC is the most potent β-lactamase in the *Mtb* proteome and is well characterized to bind and hydrolyze a broad range of β-lactam antibiotics[27]. The PBPs DacB and Rv1367c also have β-lactamase activity[42].

All enzymes were expressed and purified under native conditions (**Figure S2**). We assessed meropenem binding using an *in vitro* competition assay. Enzymes were incubated with Mero-sCy5 with or without pre-treatment with excess meropenem. Mero-sCy5 labeled all enzymes, and pretreatment with meropenem significantly reduced labeling of all enzymes except DapE (**Figure 4A,B**). Differences in fluorescent labeling of enzymes are likely due to variable β-lactam affinity, labeling kinetics, and/or the balance between inhibition and hydrolysis. Mero-sCy5 labeling, as well as its competition by meropenem, was greatest for Rv1723, Rv0309, and the positive controls BlaC and DacB. This competition assay confirmed that Rv1723, Rv2257c, DapE, Rv0309, MurI, and LipD bind meropenem, revealing these are previously undiscovered β-lactam targets in *Mtb*.

**Figure 4.**
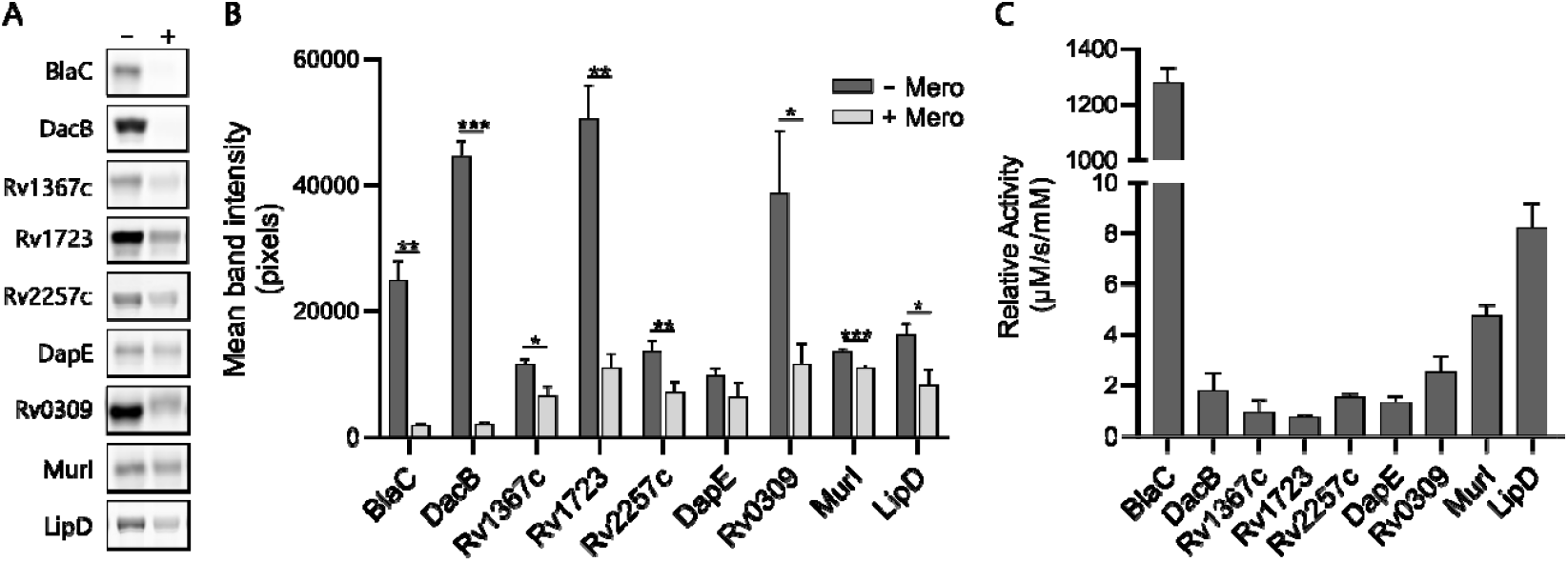
Mycobacterial enzymes identified by ABPP bind to Mero-sCy5 and exhibit β-lactamase activity. **A**. Fluorescent gel images of Mero-sCy5-labeled enzymes. Purified enzymes (2 μg) were labeled with Mero-sCy5 (1 hr) with (+) or without (−) pre-treatment with 200-fold molar excess meropenem (1 hr) and analyzed by SDS-PAGE. **B**. Quantification of signal from Mero-sCy5 labeled enzymes. Mean band intensity (n = 3) of each enzyme condition was quantified. The difference between ± Mero pretreatment was assessed by t-test (*, p ≤ 0.05; **, p ≤ 0.01; ***, p ≤ 0.001). **C**. Relative activity for each enzyme was determined by dividing the rate of nitrocefin hydrolysis by the enzyme concentration (n = 3). The rate was measured from 0-10 min with 200 μM nitrocefin for all enzymes except Rv0309 (10-20 min), which displayed a lag before acceleration of hydrolysis.

We next assessed β-lactamase activity using nitrocefin, a commonly used chromogenic cephalosporin. This assay demonstrated that all six enzymes can bind and hydrolyze the cephalosporin class of β-lactams (**Figure 4C**). BlaC rapidly hydrolyzed this substrate at a much faster rate than the other enzymes. After BlaC, the highest hydrolytic activity was observed for MurI and LipD. The other four enzymes (Rv1723, Rv2257c, DapE, and Rv0309) showed nitrocefin hydrolysis rates similar to the controls DacB and Rv1367c. The β-lactamase activity of LipD was previously investigated when it was characterized as a stress-induced lipase[108]. No β-lactamase activity was found; however, the study used denatured and refolded LipD, which may have altered enzymatic activity. While our finding contradicts that result, it is consistent with reports of other *Mtb* lipases having β-lactamase activity[65, 66].

To summarize, we validated meropenem binding and β-lactam hydrolysis by nine *Mtb* enzymes, including six with no prior evidence of β-lactam interaction. We summarize our findings in **Table 1**, which is a comprehensive list of the 32 *Mtb* proteins confirmed to date to bind β-lactam antibiotics. **Table 1** includes previously identified targets and the top hits identified by ABPP and featured in **Figure 2** and **Figure** 3. There are likely additional valid targets within our ABPP results, a possibility highlighted by our validation of meropenem binding by LipD and MurI. This highly promiscuous target profile definitively demonstrates that β-lactams bind many more proteins in *Mtb* than historically accepted and suggests a high degree of polypharmacology.

**Table 1.**
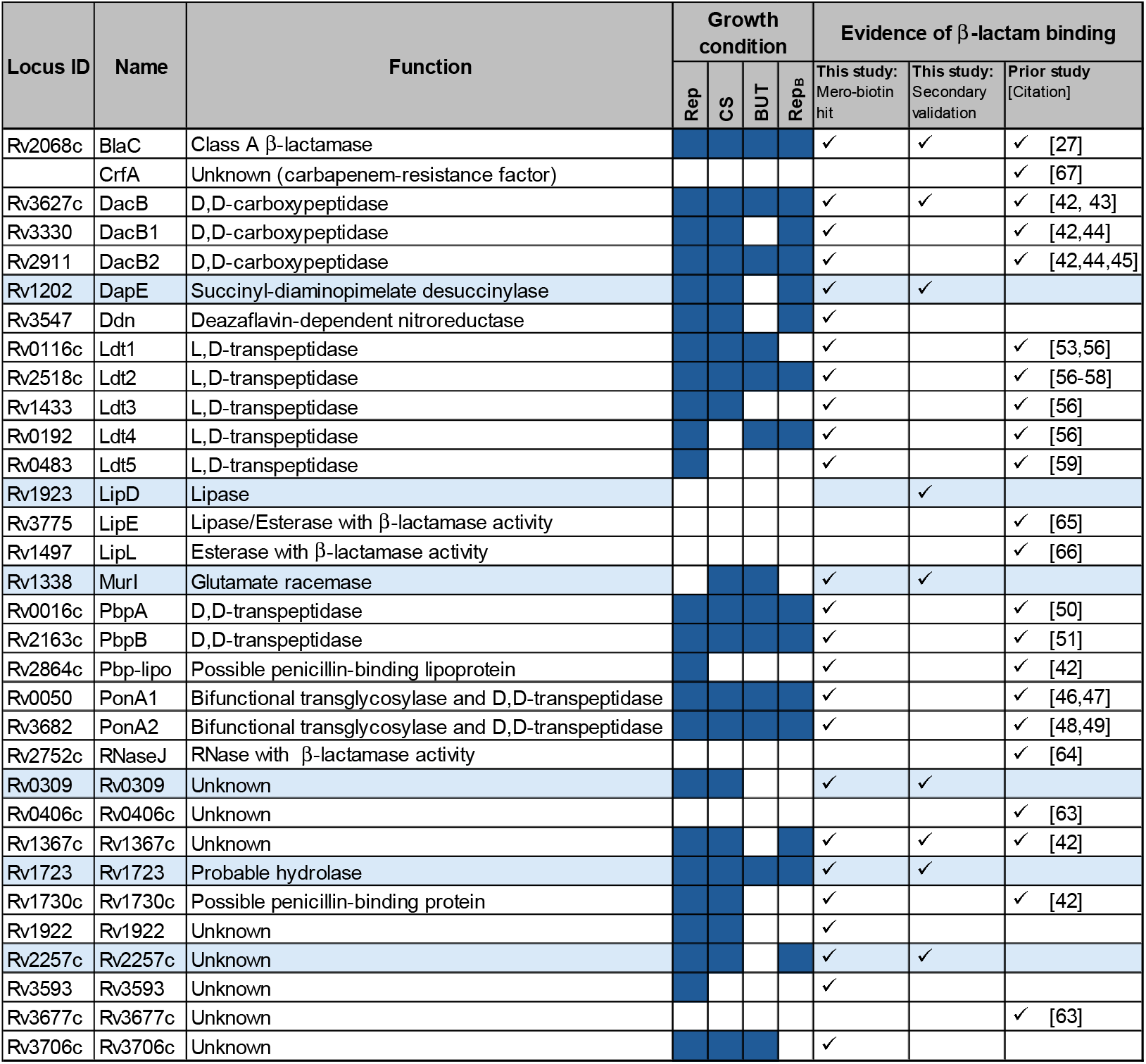
*M. tuberculosis* proteins that bind to β-lactam antibiotics.

## Kinetic characterization of Rv1723, Rv2257c, and Rv0309 β-lactamase activity

Considering that Rv1723, Rv0309, and Rv2257c displayed the highest degree of Mero-sCy5 binding of the novel targets, we further investigated the β-lactamase activity these enzymes through kinetic characterization of nitrocefin hydrolysis (**Figure 5**). We assumed that these enzymes followed Michaelis-Menten kinetics. However, the rate of hydrolysis by Rv0309 was initially slow before reaching a steady state (see **Figure S3**). We have not determined the mechanism underlying this “lag phase”, but it is consistent with Rv0309 undergoing slow activation via a conformational change[109]. Rv1723 had a brief lag as well, but the velocity stabilized quickly. The Michaelis constant (K_M_) was high micromolar for Rv1723 (K_M_ = 610 ± 150 μM), Rv2257c (K_M_ = 1000 ± 200μM), and Rv0309 (K_M_ = 740 ± 150 μM). These enzymes therefore displayed much lower affinity for nitrocefin than BlaC, which has a reported K_M_ of 60 μM[27]. The nitrocefin hydrolysis rate (V_max_) and catalytic efficiency (*k*_cat_/K_M_) of all three enzymes were also poor. These data suggest that cephalosporins are not the preferred β-lactam substrates of Rv1723, Rv0309, or Rv2257c.

**Figure 5.**
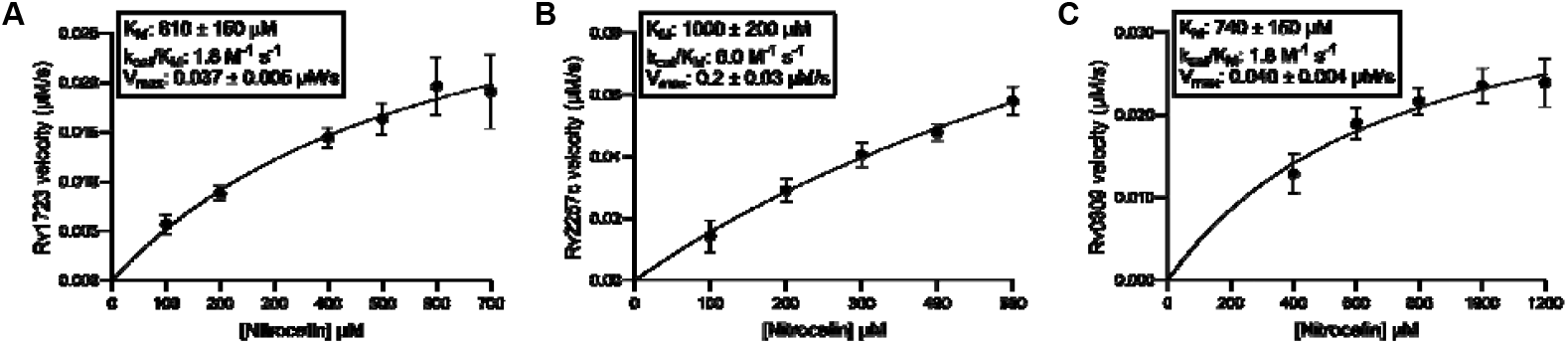
Kinetics of nitrocefin hydrolysis by Rv1723, Rv2257c, and Rv0309. The initial rate of hydrolysis versus nitrocefin concentration for Rv1723 (**A**), Rv2257c (**B**), and Rv0309 (**C**). Formation of product was monitored continuously at 37 °C in Tris buffer (pH 7.2). Kinetic values (K_M_, k_cat_/K_M_ and V_max_) were determined from at least three independent measurements and at least five different nitrocefin concentrations. Mean velocities were modeled by non-linear regression assuming Michaelis-Menten kinetics. Error bars represent the standard deviation of mean velocities.

### Assessment of Rv1723 and Rv0309 β-lactam substrate preferences

Since the kinetic analysis suggested that Rv1723 and Rv0309 are poor cephalosporinases, we sought to determine the subclasses of β-lactams that would bind favorably to these enzymes. First, we completed a competition assay with a range of β-lactams from the carbapenem (meropenem, imipenem, tebipenem), cepham (ceftriaxone, cefoxitin), penam (penicillin G, amoxicillin), and monobactam (aztreonam) subclasses. Rv1723 and Rv0309 showed distinct inhibition profiles. Labeling of Rv1723 by Mero-sCy5 was competed by pretreatment with carbapenems alone (**Figure 6A**), indicating a preference of Rv1723 for the carbapenem subclass similar to that reported for LDTs[56, 58]. Rv0309 exhibited preferential binding to carbapenems followed by amoxicillin, whereas it did not bind to the cephalosporins (**Figure 6B**). We confirmed that each enzyme’s predicted active site nucleophile was required for covalently binding meropenem by comparing Mero-sCy5 labeling of the wild-type enzyme versus an active-site mutant (Rv1723-Ser107Ala and Rv0309-Cys193Gly). The mutants showed almost no binding to Mero-sCy5 (**Figure 6A,B**). These results confirmed the identity of the active site nucleophile for both enzymes and that Mero-sCy5 labeling is dependent on activity.

**Figure 6.**
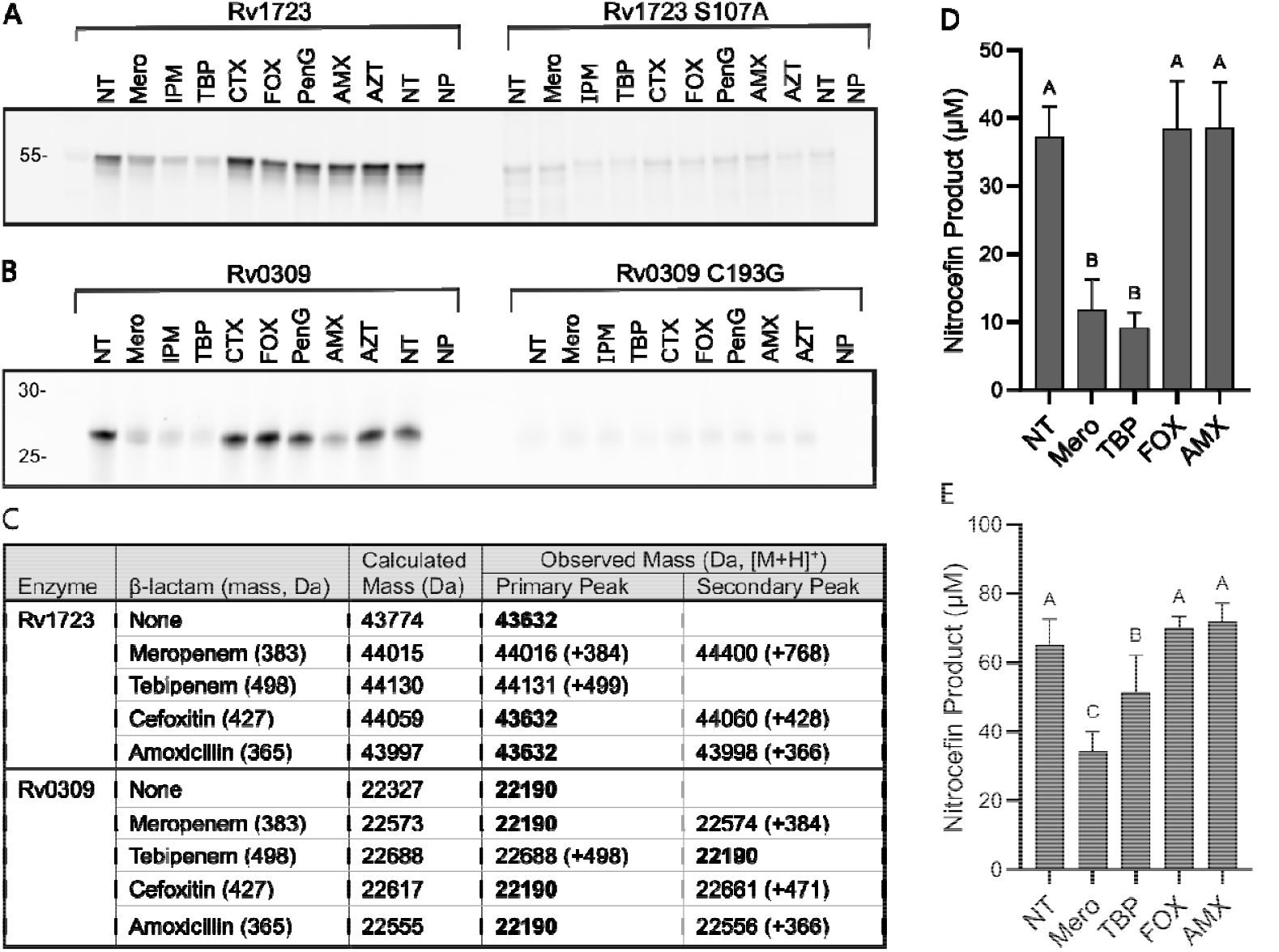
Carbapenems inhibit Rv1723 and Rv0309 more potently than other β-lactam classes. Purified Rv1723 (**A**, 2 μM) and Rv0309 (**B**, 12 μM) were treated with β-lactams (500 μM) prior to labeling with Mero-sCy5. Active site mutant enzymes (Rv1723 S107A, Rv0309 C193G) were included for comparison. Abbreviations: no treatment (NT), meropenem (Mero), imipenem (IPM), tebipenem (TBP), ceftriaxone (CTX), cefoxitin (FOX), penicillin G (PenG), amoxicillin (AMX), aztreonam (AZT), and NT + no probe (NP). **C**. Native MS analysis of Rv1723 and Rv0309 binding with β-lactams. Enzyme-only masses are presented in bold. Observed masses are presented as [M+H]^+^, with adduct masses listed in parentheses. **D, E**. Measurement of β-lactam inhibition of enzyme β-lactamase activity. Total hydrolyzed nitrocefin was measured after 2-hour incubation with either Rv1723 (**D**) or Rv0309 (**E**). Enzymes were pre-treated with β-lactams (5-fold molar excess; 30 min) as indicated. The mean product (n = 9) for each drug treatment was analyzed by ANOVA. Error bars represent standard error of the mean.

We used native mass spectrometry to confirm the results of the competition assay. Rv1723 and Rv0309 were incubated with 25-fold molar excess antibiotic before analysis (**Figure 6C**). For Rv1723, we observed a mass consistent with enzyme-acyl formation for meropenem, tebipenem, and cefoxitin. Binding to meropenem and tebipenem was complete (no unbound Rv1723 observed), while binding to cefoxitin was only partial. Rv1723 did not bind to amoxicillin. Rv0309 bound tebipenem almost completely (negligible unbound enzyme observed). Rv0309 partially bound meropenem and displayed surprisingly high binding to amoxicillin (~50% bound versus unbound enzyme). No notable binding to cefoxitin was observed. These native mass spectrometry results are consistent with our competition assay. The interaction between Rv0309 and amoxicillin observed across these assays was unexpected given that penams generally do not bind the other *Mtb* LDTs. Structural analysis of Rv0309 is warranted to determine how this enzyme discriminates between β-lactams.

Lastly, we assessed nitrocefin hydrolysis by these enzymes in the presence of β-lactams (**Figure 6D,E**). Despite these enzymes being poor cephalosporinases, we wanted to assess whether binding of preferred β-lactams affected enzymatic activity. Rv1723 and Rv0309 were incubated with a 5-fold molar excess of antibiotic before measuring nitrocefin hydrolysis. Meropenem and tebipenem significantly reduced the amount of nitrocefin hydrolyzed by both enzymes, while amoxicillin and cefoxitin did not. This result further supports the preference of Rv1723 and Rv0309 for binding to carbapenems and demonstrates that their β-lactamase activity is inhibited by meropenem.

## Conclusions

Emerging clinical and experimental evidence suggests high, but unrealized, potential of β-lactam antibiotics for treating TB[18-21]. Progress in this area has been hindered by an incomplete understanding of the mechanistic basis of β-lactam activity against *Mtb*. More specifically, there was no complete list of enzymes inhibited by this class of antibiotics. For decades, it was thought that β-lactams only inhibited *Mtb*’s PBPs[40], but this view is incorrect. In the current study, we provide comprehensive evidence that many other enzymes are targeted by meropenem. Using a new ABP, Mero-biotin, we have stringently identified a diverse set of targets, including PBPs, LDTs, β-lactamases, and other functionally distinct enzymes. We provide validation of six novel β-lactam targets: Rv1723, Rv2257c, DapE, Rv0309, MurI, and LipD.

Our results show that β-lactams bind not only to enzymes that crosslink PG side chains but also those associated with upstream PG metabolism, such as MurI and DapE. MurI and DapE have been structurally and biochemically characterized[98, 106], and their interactions with β-lactams were unexpected based on these reports. To our knowledge, we present the first evidence of a glutamate racemase interacting with a β-lactam drug. We are captivated by the possibility that MurI inhibition contributes to the β-lactam mechanism of action in *Mtb*. We plan to further evaluate the interaction of β-lactam antibiotics with MurI.

We propose that at least 32 *Mtb* enzymes bind β-lactams, and these interactions collectively contribute to the potency of carbapenems against *Mtb*. Most of these enzymes were active and inhibited by meropenem under physiologically relevant conditions representative of both acute and chronic TB. We hypothesize that pronounced polypharmacology underlies the efficacy of β-lactams as a class of drug. Prior studies and the current work support this idea in *Mtb*. It is possible that related pathogens, including *Mycobacterium abscessus*, have similarly large numbers of enzymes inhibited by β-lactams. It is worth considering if antibiotic drug development should focus less on identifying highly specific drugs in favor of a promiscuous mechanism of action that is tolerated by patients.

## Supporting information

Electronic Supporting Information

## Acknowledgements

Funding for this research was provided by the National Institute of Health (NIAID: R01 AI149737). Pacific Northwest National Laboratory is operated by Battelle for the Department of Energy (DOE) under contract DE-AC05-76RL01830. A portion of this research was supported by an Environmental Molecular Sciences Laboratory (EMSL) user project award (https://www.emsl.pnnl.gov/project/60231) for leveraging instrument capabilities operating at EMSL, a DOE Office of Science User Facility. Development of chemoprotR was supported by an EMSL Science and Technology Developer project award (https://www.emsl.pnnl.gov/project/60473). Kurt Krause (University of Otago) generously provided the plasmid encoding MurI. We thank Dr. Jordan Devereux (OHSU Medicinal Chemistry Core; RRID: SCR_019048) for synthesizing Mero-sCy5 and Prof. Ujwal Shinde (OHSU Biophysics Core; RRID: SCR_022744) for assistance purifying Rv1723. We are grateful to Priscila Lalli, Vanessa Paurus, and Ron Moore for proteomics runs.

## Abbreviated Methods

Detailed descriptions of protocols are provided in the ESI. This includes synthetic methods, *Mtb* culture conditions, proteomics pipelines, enzyme preparation, and enzyme analysis.

### Mycobacterial Culture Conditions

*Mycobacterium tuberculosis* (*Mtb*) mc^2^6020 (Δ*lysA*, Δ*panCD* mutant)[110] was cultured in 7H9-OADC media supplemented with lysine, pantothenate, and casamino acids. Carbon starvation was induced by growth in bottles (5 weeks) in minimal media, as described[111]. Alternatively, a lipid-rich condition was modeled by growing *Mtb* in MOPS-buffered media supplemented with butyrate, as described[91]. Six biological replicates were grown for each condition.

### LC-MS/MS Proteomics analysis

Lysates were prepared by mechanical disruption of cells in PBS supplemented with 0.5% n-dodecyl-D-β-maltoside. Sterilized lysates were quantified by BCA assay and normalized for global proteomics analysis (100 μg total protein, n = 6 per condition). For ABPP, lysates were labeled with 10-30 μM Mero-biotin (600 μg total protein, n = 6 per condition) or vehicle (no probe controls [NPC], n = 4) for 60 min (room temperature). Excess probe was removed and samples were affinity-enriched on streptavidin-agarose resin with extensive washing. On-bead trypsin digestion yielded peptides for LC-MS/MS analysis.

Enriched peptides were analyzed using a Waters nanoAcquity ultra performance liquid chromatography (UPLC) system connected to a Q Exactive Plus Orbitrap mass spectrometer with data dependent acquisition (Thermo Scientific, San Jose, CA). Separations were performed using C18 chromatography with a gradient of 1-75% mobile phase B (acetonitrile + 0.1% formic acid) in mobile phase A (0.1% formic acid in water). Additional details of the UPLC and mass spectrometer settings are described in the ESI. Proteomics data analysis was performed with MASIC[112], MSGF+[113], and Mage[114] software using the *M. tuberculosis* H37Rv protein database (UniProt taxon ID 83332, downloaded on 2021-03-07). Additional processing was performed using ChemoprotR, a custom R script developed at PNNL; detailed data processing methods are described in the ESI.

### Enzyme expression and purification

Constructs were previously described (BlaC, DacB, Rv1367c, and MurI) or designed and purchased (Genscript). Enzymes were expressed and purified as described[42]. Briefly, protein expression in *E. coli* (BL21-star-DE3) was induced overnight. Cells were lysed by sonication and soluble proteins were purified by immobilized metal affinity chromatography (i.e., Ni-NTA agarose). Enzyme purity was assessed by SDS-PAGE analysis and most enzymes were used without further purification.

### Enzyme labeling and fluorescent gel analysis

Lysates or purified enzyme were diluted in buffer to normalize total protein concentration. For samples pre-treated with antibiotics, ~100-fold excess antibiotic was added 1 hr prior to probe labeling. Samples were labeled with Mero-sCy5 (5-10 μM) for 1 hr at RT. Samples were denatured, resolved by SDS-PAGE, and imaged on a Typhoon multimode imager (Cy5: λex/λem = 635/670 BP 30 nm). Images were processed and analyzed in ImageJ[115].

### Nitrocefin hydrolysis assays

The hydrolysis of nitrocefin was quantified by measuring the formation of product at 486 nm (ε = 20 500 M^−1^ cm^−1^). Reactions were initiated by combining enzyme with nitrocefin in 50 mM Tris-Cl (pH 7.2 at 37 °C), 150 mM NaCl. Data from three independent replicates were analyzed in GraphPad Prism (v. 10.2.0).

