## Supplementary material for "Comprehensive identification of β-lactam antibiotic polypharmacology in *Mycobacterium tuberculosis*": Electronic Supporting Information

**Table of Contents**

**Materials and Methods**

**I. Chemicals	2**

**II. Synthesis of ABPs: Meropenem-biotin and Meropenem-sCy5	2**

**III. Activity-based protein profiling	3**

Mycobacterial Culture	3

*Mtb* lysis	4

Labeling and Enrichment of Lysates for ABPP	4

Mass Spectrometry Sample Preparation	5

Liquid chromatography tandem MS	5

Analysis of MS data	5

Western blot analysis	6

**IV. Enzyme hit validation	6**

Genetic construction of plasmids	6

Small-scale expression and purification	6

Large-scale expression and purification	6

In-Gel Analysis of Mero-sCy5 Binding and Competition	7

Relative Activity of Mtb Enzymes	7

Kinetics analysis: Rv1723, Rv2257c, and Rv0309	7

Inhibition of Activity by β-lactams	8

Native Mass Spectrometry of Enzyme-Inhibitor Complexes	8

**Supplementary Data**

Schemes S1 and S2	9-10

Figures S1-S6	11-15

**Supplementary Tables**

Table S1: *Mtb* proteins with endogenous biotinylation or biotin binding.	16

Table S2: *See corresponding Excel file “ESI Table S2_ABPP hits”.*

Table S3: *See corresponding Excel file “ESI Table S3_ABPP hit overlap”.*

Table S4: *M. tuberculosis* mc^2^6020 Culture Media Recipes.	17

Table S5: Summary of *Mtb* “hit” protein constructs.	18

Table S6: Primers for Site-directed mutagenesis (SDM)	19

**Materials and Methods**

**I. Chemicals**

Unless otherwise noted, all chemicals were purchased from Sigma-Aldrich, ThermoFisher Scientific, or Lumiprobe and used as received. Antibiotics used in the present study were sourced as follows: meropenem (AA Blocks, CAS 1192500-31-4), imipenem (Thermo Scientific, CAS 74431-23-5), tebipenem (AA Blocks, CAS 161715-24-8), ceftriaxone (Acros Organics, CAS 104376-79-6), cefoxitin (Sigma-Aldrich, CAS 33564-30-6), penicillin G (Thermo Scientific, CAS 69-57-8), amoxicillin (Alfa Aesar Chemicals, CAS 61336-70-7), aztreonam (AA Blocks, CAS 78110-38-0). Structures of antibiotics are provided in **Scheme S2**. Antibiotic stock solutions were prepared fresh in water or DMSO (tebipenem) immediately prior to use.

**II. Synthesis of ABPs: Meropenem-biotin and Meropenem-sulfoCy5**

Unless otherwise stated, reactions were magnetically stirred in flame-dried glassware under an atmosphere of nitrogen. Anhydrous solvents were purchased in septum-sealed bottles and stored under nitrogen. Semi-dry DMF was stored over 4 Å molecule sieves. Et_3_N was stored over K_2_CO_3_ in a desiccator. Reactions were monitored by thin-layer chromatography (TLC) on glass-backed silica gel plates (Silicycle 60 Å, 250 µM). Reverse phase column chromatography was performed with the indicated solvents on a Biotage Isolera One automated chromatography system.

Mass spectra were acquired at Portland State University's bioanalytical mass-spectrometry facility on a ThermoElectron LTQ-Orbitrap Discovery high-resolution mass spectrometer with electrospray ionization (ESI-HRMS).

^1^H-NMR spectra were taken at ambient temperature in the indicated solvent at Portland State University's NMR facility on a Bruker Avance II at 400 MHz or at OHSU (Department of Chemical Physiology & Biochemistry) on a Bruker Avance Neo NanoBay at 400 MHz. Spectra were calibrated to the residual solvent peak. Chemical shifts are reported in ppm. Coupling constants (*J*) are reported in Hertz (Hz) and rounded to the nearest 0.1 Hz. Multiplicities are defined as: s = singlet, d = doublet, dd = doublet of doublets, dt = doublet of triplets, t = triplet, td = triplet of doublets, quin = quintet, m = multiplet, br s = broad singlet.

**Generation of Biotin Acid Chloride**

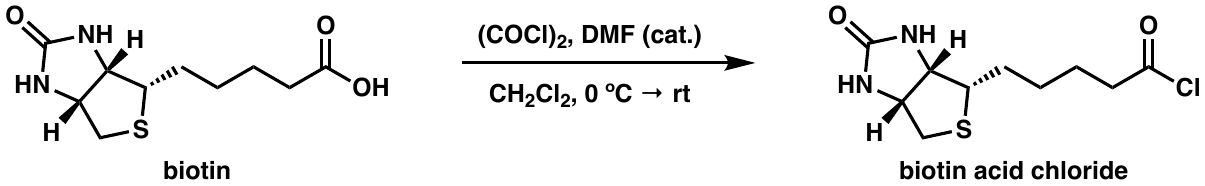
In a 5 mL collection flask (dry, under N_2_), biotin (0.2447 g, 1.00 mmol, 1 equiv, Ark Pharm AK-44010) was combined with oxalyl chloride (0.176 mL, 2.05 mmol, 2.05 equiv, Acros Organics 129610250) and CH_2_Cl_2_ (1.5 mL, 0.67 M, Aldrich 270997-1L) to give a white suspension, which was cooled to 0 °C. A catalytic drop of dry DMF was added. After stirring for 20 min at 0 °C the flask was allowed to warm and stirred at RT for 1 h. The resulting white powder was left to dry *in vacuo* overnight and used without further purification.

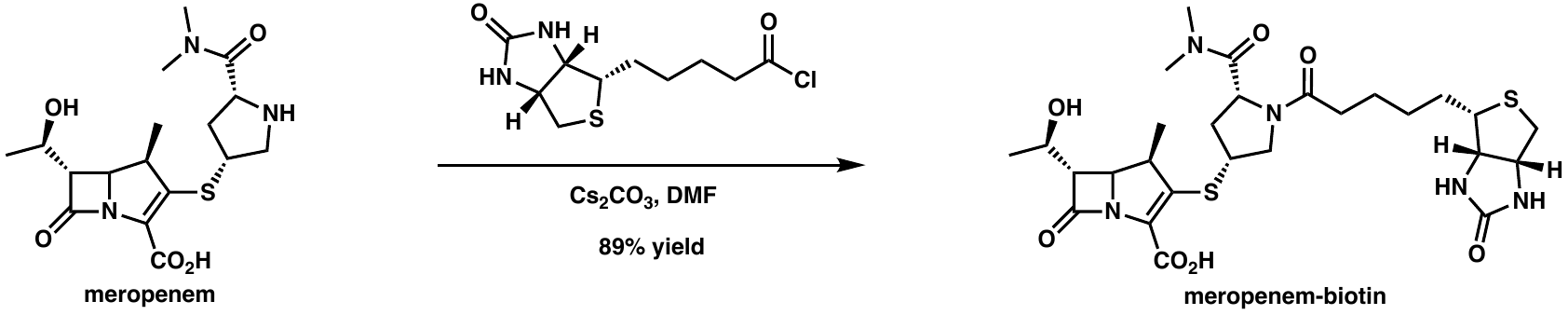
**Synthesis of Meropenem-biotin**

Meropenem trihydrate (0.0443 g, 0.101 mmol, 1 equiv, Carbosynth) was dissolved in semi-dry DMF (0.5 mL) in a 2 dram vial. Cs_2_CO_3_ (0.098 g, 0.304 mmol, 3 equiv) was added and the suspension stirred at RT. Biotin acid chloride (0.0398 g, 0.152 mmol, 1.5 equiv) was added in one portion and the resulting heterogeneous mixture stirred at RT. After 40 min, additional Cs_2_CO_3_ (0.098 g, 0.304 mmol, 3 equiv), biotin acid chloride (0.0398 g, 0.152 mmol, 1.5 equiv), and DMF (0.25 mL) were added. After 75 min additional biotin acid chloride (0.0398 g, 0.152 mmol, 1.5 equiv) was added. After 95 min, the reaction mixture was transferred to a microcentrifuge tube and the Cs_2_CO_3_ pelleted. The pellet was resuspended in 0.5 mL DMF and centrifuged again. This step was repeated. The DMF layers (~2 mL) were combined and triturated with Et_2_O (~13 mL). After centrifugation, the pellet was resuspended via vortexing and sonication in Et_2_O (2 x 15 mL), and then dried under vacuum to give an off-white solid (0.1622 g, 263% yield), which retained some DMF and/or H_2_O. This compound was further purified by reverse phase chromatography on a Biotage Isolera system with a SNAP KP-C18-HS cartridge. The method used was 5% CH_3_CN in 95% H_2_O (both doped with 0.1% formic acid) for 1 column volume, followed by a gradient of 5-50% of the same solvent system over 10 column volumes. The fractions containing the product were combined and dried by lyophilization to give a fluffy off-white powder (0.0547 g, 89% yield). The ESI-HRMS [M+H]^+^ m/z calculated for meropenem-biotin: 610.23; observed: 610.235.

^1^H-NMR (400 MHz; D_2_O): δ 4.89 (m, 1H), 4.62 (dd, *J* = 7.8, 5.0 Hz, 1H), 4.44 (dd, *J* = 7.9, 4.5 Hz, 1H), 4.26 (ddt, *J* = 22.0, 10.6, 4.9 Hz, 3H), 3.98 – 3.87 (m, 1H), 3.60 – 3.52 (m, 2H), 3.50 (dd, *J* = 6.1, 2.5 Hz, 1H), 3.40 – 3.31 (m, 1H), 3.20 – 3.12 (m, 1H), 3.16 (s, 3H), 2.96 (s, 3H), 2.92 – 2.83 (m, 1H), 2.79 (d, *J* = 13.1 Hz, 1H), 2.43 (t, *J* = 6.9 Hz, 2H), 1.89 – 1.79 (m, 1H), 1.74 (dt, *J* = 13.6, 6.6 Hz, 1H), 1.62 (m, 3H), 1.51 – 1.40 (m, 2H), 1.31 (d, *J* = 6.3 Hz, 3H), 1.26 (d, *J* = 6.9 Hz, 3H). Spectra are provided in **Figure S4**.

**Synthesis of Meropenem-sulfoCy5**

Meropenem-sulfoCy5 (Mero-sCy5) was synthesized by the OHSU Medicinal Chemistry core following a published protocol[1]. The compound was purified by HPLC chromatography to 85% purity. The ESI-HRMS [M+H]^+^ m/z calculated for Mero-sCy5: 1008.2; observed 1008.46. Mero-sCy5 was observed to be unstable in some solvents (e.g., DMSO), which prevented structural analysis by NMR. Single use aliquots were prepared (2 mM in 10 mM HEPES pH 7.5) and stored at -80 °C until use.

**III. Activity-based protein profiling**

**Mycobacterial Culture Conditions**

*Mycobacterium tuberculosis* (*Mtb*) mc^2^6020 (Δ*lysA*, Δ*panCD* mutant)[2], a double auxotrophic mutant derived from the laboratory strain *Mtb* H37Rv, was obtained from W. Jacobs laboratory (Albert Einstein College of Medicine and HHMI) and handled as a BSL-2 pathogen.

Bacteria were thawed from frozen stocks stored at -80 °C in 30% glycerol. *Mtb* were cultured in 7H9/OADC-KPC medium (**Table S4**). Cultures were grown at 37 °C with 100 rpm in aerated polycarbonate shake flasks with a 0.2 μm filter cap (Weaton #WPFPC0500S). For proteomics, six replicates of carbon starvation and butyrate-rich cultures, along with separate sets of replicating cultures matched to each, were grown as specified below. All cultures were harvested at specified time points (5 min, 4000 xg, 4 °C), washed twice with PBS, and stored at -30 °C in PBS until lysis.

*Carbon Starvation and Matched Replicating Cultures*

Culture conditions to induce dormancy via carbon starvation were as described[3]. Briefly, *Mtb* were grown to an OD_600_ of 0.8 - 1.2, washed, and diluted (OD_600_ of 0.2) in carbon starvation medium (7H9/Tx-KP, **Table S4**). Cultures (CS, n=6, 300 mL) were grown standing at 37 °C in 1 L plug-sealed bottles (Corning #430195) for 5 weeks. Matched replicating cultures were simultaneously prepared from the same washed cell stock after dilution (OD_600_ of 0.2) in 7H9/OADC-KPC medium. Cultures (Rep, n=6, 200 mL) were grown shaking (100 rpm, 37 °C) in aerated 500 mL shake flasks until an OD_600_ of ~1.0 was reached. Cells were harvested through centrifugation (5 min, 4000 xg, 4 °C), washed twice with PBS, and stored at -30 °C in PBS until lysis.

*Butyrate-Rich and Matched Buffered Replicating Cultures*

Culture conditions to mimic the lipid-rich environment of TB granulomas were adapted from previously reported methods[4]. *Mtb* were grown in 7H9/OADC-KPC to an OD_600_ of 0.8 - 1.2. Cells were acclimated to butyrate-rich conditions through two cycles of dilution and growth as follows. Cells were washed twice in butyrate-rich medium (7H9/Butyrate-KP **Table S4**). Washed cells were diluted to an OD_600_ of 0.05 in 7H9/Butyrate-KP and grown to an OD_600_ of 0.8 – 1.2. After acclimation, cells were diluted to an OD_600_ of 0.05 and cultures (Butyrate, n=6, 300 mL) were grown shaking (100 rpm, 37 °C) in aerated 500 mL shake flasks until an OD_600_ of 0.8-1.2 was reached. Matched buffered replicating cultures were simultaneously prepared from the same stock cultures. Cells were washed twice with 7H9/OADC/MOPS-KP (**Table S4**). Cells were diluted to an OD_600_ of 0.05 in 7H9/OADC/MOPS-KP. Cultures (Rep_B_, n=6, 300 mL) were grown shaking (100 rpm, 37 °C) in aerated 500 mL shake flasks until an OD_600_ of 0.8-1.2 was reached.

**Preparation of *Mtb* Lysates**

Cells were lysed as previously described [3]. Briefly, whole cell lysates were obtained by mechanical disruption in 0.5% n-dodecyl-D­-β-maltoside (Chem-Impex #21950, CAS 69227-93-6) in PBS (PBS-DM). Lysates were filtered twice through 0.2 µm PVDF membrane filters (13 mm, Pall) to sterilize. A bicinchoninic acid (BCA) assay (Pierce) was used to quantify the total protein concentration of all lysates.

**Labeling and Enrichment of Lysates for ABPP**

A modified protocol based on the methods of Brandvold et al.[5] was optimized for affinity enrichment of meropenem-biotin-labeled *Mtb* lysates. Samples for global proteomics analysis (n=6 per culture condition) were also prepared alongside ABPP samples. Mero-biotin-probed and no probe control samples were analyzed by anti-biotin western blot to preliminarily assess labeling and sample quality (**Figure S5**). All samples were stored at -20 °C before transfer to Pacific Northwest National Laboratory (PNNL) for LC-MS/MS analysis.

*Carbon Starvation and Matched Replicating Samples*

Lysates (600 μg total protein) were incubated with either 30 μM meropenem-biotin or vehicle (no probe controls [NPC], n=4) (60 min, RT). Protein was purified of excess probe via 3 kDa molecular weight cutoff (MWCO) filtration (Amicon). Biotinylated proteins were affinity purified on streptavidin-agarose resin (Thermo Fisher Scientific, 20353). Samples were bound to prewashed resin in 1% SDS in PBS at a protein:settled-resin ratio (μg:μL) of 4:1 (60 min, RT, end-over-end rotation). Resin-bound protein was washed within fritted chromatography columns (Bio-Rad, 7326008) as follows: 1x 1% SDS in PBS, 3x 0.5% SDS in PBS, 1x 6 M urea in 25 mM ammonium bicarbonate, 3x ultrapure water, 4x PBS, 4x 25 mM ammonium bicarbonate, pH 8. Resin-bound protein was then trypsin (Promega) digested at a trypsin:protein ratio (μg:μg) of 1:4000 (overnight, RT, end-over-end rotation). Digested peptides were separated from resin (2000 xg, 5 min, RT) and residual peptides were collected off resin with additional 25 mM ammonium bicarbonate (30 min, RT, end-over-end rotation).

*Butyrate-Rich and Buffered Replicating Samples*

Lysates (600 μg total protein) were incubated with either 10 μM meropenem-biotin or vehicle (NPC, n=3) (60 min, RT). Protein was purified of excess probe via methanol precipitation (10-fold excess, -80 °C, 30 min). Precipitated protein was pelleted (4800 xg, 20 min, 4 °C) and air-dried before resuspension in 1% SDS in PBS. Biotinylated proteins were affinity purified on streptavidin-agarose resin (Thermo Fisher Scientific, 20353). Samples were bound to prewashed resin in 1% SDS in PBS at a protein:settled-resin ratio (μg:μL) of 4:1 (60 min, RT, end-over-end rotation). Resin-bound protein was washed within fritted chromatography columns (Bio-Rad, 7326008) as follows: 1x 1% SDS in PBS, 2x 0.5% SDS in PBS, 2x ultrapure water, 2x PBS, 2x 25 mM ammonium bicarbonate, pH 8. Resin-bound protein was then trypsin (Promega, V511A) digested at a trypsin:protein ratio (μg:μg) of 1:4000 (overnight, RT, end-over-end rotation). Digested peptides were separated from resin (2000 xg, 5 min, RT) and residual peptides were collected off resin with additional 25 mM ammonium bicarbonate (30 min, RT, end-over-end rotation).

**Mass Spectrometry Sample Preparation**

Peptides were evaporated to dryness in a SpeedVac concentrator, and then dried peptides were reconstituted in 40 µL of 25 mM ammonium bicarbonate and heated at 37 °C for 5 min at 1000 rpm on a thermoshaker. Samples were briefed centrifuged to pellet any insoluble debris, and the samples transferred to ultracentrifuge tubes. Samples were ultracentrifuged at 53,000 rpm in a Beckman ultracentrifuge for 20 min at 4 °C. After centrifugation, 25 µL of supernatant was carefully transferred to LC-MS vials and stored at -20 °C until ready for LC-MS/MS analysis.

**Liquid Chromatography Tandem Mass Spectrometry**

ABPP samples were analyzed using a Waters nanoAcquity ultra performance liquid chromatography (UPLC) system connected to a Q Exactive Plus Orbitrap mass spectrometer (Thermo Scientific). Samples were loaded into a precolumn (150 μm i.d., 4 cm length, packed in-lab with Jupiter C18 packing material, 300 Å pore size, 5 μm particle size; Phenomenex) using mobile phase A (0.1% formic acid in water). The separation was carried out in a LC column (Packed in-lab into an empty self pack NanoLC column (CoAnn Technologies) 75 µm i.d., 30-cm column with Waters BEH C18 packing material, 130-Å pore size, 1.7 µm particle size (Waters Corporation)) at a flow rate of 200 nL/min using a 60 min gradient of 1-75% mobile phase B (acetonitrile + 0.1% formic acid). To prevent carryover, the column was washed with 95-50% mobile phase B for 20 min and equilibrated with 1% mobile phase B for 30 min before the next sample injection. The mass spectrometer source was set at 2.2 kV, and the ion transfer capillary was heated to 300 °C. The data-dependent acquisition mode was employed to automatically trigger the precursor scan and the MS/MS scans. The MS1 spectra were collected at a scan range of 300-1800 m/z, a resolution of 70,000, an automatic gain control (AGC) target of 3 x 10^6^, and a maximum injection ion injection time of 20 ms. For MS2, top 12 most intense precursors were isolated with a window of 1.5 m/z and fragmented by higher-energy collisional dissociation (HCD) with a normalized collision energy at 30%. The Orbitrap was used to collect MS/MS spectra at a resolution of 17,500, a maximum AGC target of 10^5^, and maximum ion injection time of 50 ms. Each parent ion was fragmented once before being dynamically excluded for 30 s.

**Analysis of Mass Spectrometry Data**

MS/MS Automated Selected Ion Chromatogram generator (MASIC) was used to generate selected ion chromatograms (SICs) for all of the parent ions chosen for fragmentation in the LC-MS/MS data[6]. MSGF+ software[7] was then used to perform peptide searches against the *M. tuberculosis* H37Rv protein database (UniProt taxon ID 83332, downloaded on 2021-03-07) with the following parameters: parent ion tolerance of 20 ppm; methionine oxidation (+15.9949 Da) as a dynamic modification. Mage software (Version 1.5.8987, <https://github.com/PNNL-Comp-Mass-Spec/Mage>) was used to extract the MSGF+ first hits results and export data for further processing.

ABPP analyses were conducted using chemoprotR, an R package currently available to EMSL users via GitLab (<https://code.emsl.pnl.gov/multiomics-analyses/chemoproteomicsR/-/tree/main>). Data were filtered such that only peptides with an MSGF spectral probability of less than 1.56 x 10^-8^ were retained, based on calculations for a 1% false discovery rate (FDR) from M. tuberculosis global proteomics data as described by Elias and Gygi[8]. Once filtered, peptide redundancies were removed by summing reporter ion masses. Reporter ion values were log2 transformed and potential outliers were calculated using a robust mahalanobis distance[9]. This distance was calculated by using multiple metrics such as the correlation of samples to other samples in the same protocol, skewness, and MAD (median absolute distance) of the molecule abundance profile, and the proportion of missing values. Data were then normalized using median centering with a group-specific backtransformation[10,11]. Peptide-level data were rolled up to a protein-level using the method, “rollup”[12]. We noted that peptide to protein rollup method may impact results and therefore special attention to the method used may be warranted[13]. Once rolled up to protein level, reverse hits and contaminants were removed from the dataset.

Statistical differences in protein intensity between ABPP groups (ABP vs. NPC) was assessed via analysis of variance (ANOVA) and independence of missingness (IMD, G-test) [14]. In cases when a protein was observed in at least two samples in each group, it was considered significantly different if the mean log2 intensity between groups had a p ≤ 0.05 by ANOVA. In cases when a protein was observed in less than two samples in one group, the number of observed values between groups was considered statistically different if P ≤ 0.05 by IMD. To identify a protein as specifically labeled by meropenem-biotin, the following criteria were applied: (1) present in at least 3 ABP replicates, (2) a difference between ABP sample and NPC mean intensity with p < 0.05 (IMD-ANOVA), (3) an at least 2-fold higher intensity in ABP samples relative to NPC samples. Proteins known to be biotinylated or biotin-binding (see **Table S1**) were manually removed from the lists.

**Western Blot Analysis**

Proteins were denatured with 1X TCEP SDS-PAGE loading dye (5X stock: 10% SDS, 289 mM Tris, 50 mM TCEP, 0.025% [w/v] Ponceau S Red, and 63% glycerol) by heating (5-10 min, 75 °C). Normalized protein (5 μg) was resolved via SDS-PAGE on a 4-12% Bis-Tris gel (Bio-Rad, 1X XT MOPS running buffer). Proteins were either transferred to a membrane for western blot analysis or stained with Coomassie G-250 (Fisher, CAS 6104-59-2). Stained gels were imaged on a flat-bed scanner (Epson).

Resolved protein for western blot analysis were transferred to a PVDF membrane (100 V, 60 min, Tris/glycine/MeOH buffer). Blots were blocked with 5% bovine serum albumin (BSA) in 1X TBST (1 M TBS, 0.1% Tween 20) (1 h, RT). Blots were washed 3x in 1X TBST and incubated with streptavidin-HRP (Thermo Scientific, S911) in 3% BSA in 1X TBST (30 min, RT). Blots were washed 3x in 1X TBST, developed with SuperSignal West Pico PLUS (Thermo Scientific, 34580), and imaged on a MyECL Imager (Thermo Scientific). The brightness and contrast of the acquired images were adjusted in ImageJ.

**IV. Enzyme Hit Validation**

**Genetic Construction of Plasmids Encoding Mycobacterial Enzymes**

Detailed information for all constructs is provided in **Table S5**. The plasmids for BlaC, DacB, Rv1367c, and MurI were previously described[15,16]. For novel enzyme hits, amino acid sequences were assessed in Uniprot and signal sequences and transmembrane portions were removed as needed. Constructs were ordered from Genscript for cloning into pET28a vector with an N-terminal histidine tag and a TEV cleavage site. It was necessary to fuse Rv2257c and LipD to a maltose-binding protein to improve solubility; these were placed into a pParallel_His7-MBP-TEV construct. We selected plasmids with kanamycin resistance because trace β-lactamase activity could interfere with subsequent assays.

Active site mutants of Rv1723 (S107A) and Rv0309 (Cys193Gly) were generated by site-directed mutagenesis (**Table S6**). Mutations were confirmed by sequence analysis.

**Small-scale Expression and Purification of Mycobacterial Enzymes**

Plasmids were transformed into *E. coli* BL21-Star-(DE3) cells [Invitrogen] to express each mycobacterial enzyme. Single colonies were used to inoculate flasks of 2XYT media (50 mL) supplemented with kanamycin and grown at 37 °C (225 rpm) until an OD_600_ of 0.4 to 0.8 was reached. Flasks were moved to 20 °C and induced with 0.25 mM IPTG overnight (20 h). Cells were pelleted by centrifugation and resuspended in 50 mM Tris-Cl (pH 8.0 at 4 °C), 150 mM NaCl, 1% sarkosyl (w/v) supplemented with 0.1 mM TCEP and 0.1 mg/mL lysozyme. After 30 min, cells were lysed by sonication on ice (40% amplitude, 20 sec on/off, 4 cycles). Lysate was clarified by centrifugation and then incubated with 750 mL Ni-NTA agarose slurry (Qiagen; pre-equilibrated with 5-10 mM imidazole). Enzymes were batch purified by washing with ~30 column volume (CV) of wash buffer (50 mM NaH_2_PO_4_, 300 mM NaCl, 20 mM imidazole, pH 8.0). Protein was eluted (3x 3CV) in 50 mM NaH_2_PO_4_, 300 mM NaCl, 250 mM imidazole, pH 8.0. Purity and size were assessed by SDS-PAGE (see **Figure S2**); the first elution appeared >95% pure. Enzyme concentrations were determined using a Pierce BCA assay kit (ThermoFisher) and ranged between 0.1 – 0.6 mg/mL. Enzymes were used directly in subsequent assays, as described, to generate data shown in **Figure 3**.

**Large-scale Expression and Purification of Rv1723, Rv2257c, and Rv0309**

Plasmids were transformed into *E. coli* BL21-Star-(DE3) cells (Invitrogen). A starter culture was used to inoculate a 2L flask of 2xYT supplemented with kanamycin (50 mg/mL). Protein expression was induced with 0.25 mM IPTG overnight (Rv1723: 16.5 hr, 20 °C; Rv2257c: 18 hr, 18 °C, Rv0309: 18 hr, 20 °C). Cells were pelleted by centrifugation and frozen before purification.

Cells were lysed in 50 mM Tris-Cl (pH 8.0 at 4 °C), 150 mM NaCl, 1% sarkosyl (w/v) by sonication. Clarified lysate (10,000 xg, 20 min, 4 °C) was bound to Ni-NTA resin (Qiagen; 4 °C, overnight). Resin was washed with: 50 mM NaH_2_PO_4_ (pH 8.0), 300 mM NaCl with 10 mM imidazole (2 CV) or 20 mM imidazole (4x 16 CV). Protein was eluted with 250 mM imidazole (4x 10 CV). All unbound, wash, and elution fractions were analyzed by SDS-PAGE. Eluted protein was concentrated and exchanged into 50 mM Tris (pH 8.0 at 4 °C), 150 mM NaCl using Amicon Ultra 10K MWCO centrifugal filters (Millipore; 4000 xg, 4 °C). Enzyme concentrations were determined by BCA assay (Pierce). Purified enzymes were stored at 4 °C (Rv1723) or at -80 °C (with 10% glycerol). We estimated our yield of each enzyme per gram (wet weight) of cell pellet as: 3 mg Rv1723; 2 mg Rv2257c; 6 mg Rv0309.

**In-Gel Analysis of Mero-sCy5 Binding and Competition**

Normalized total protein lysates or purified enzymes were labeled with Mero-sCy5 (5-10 μM, 60 min, 37 °C, protected from light). For enzyme competition experiments, samples were pretreated with β-lactam antibiotics (500 μM, 1 hr, 37 °C) prior to probe labeling. Labeling reactions were quenched by the addition of 5x TCEP loading dye. Samples were heated to 75 °C for 5−10 min and 1-5 μg of protein was resolved via SDS-PAGE (Criterion Bis-Tris gels, XT-MOPS running buffer, BioRad). Gels were de-stained of free probe overnight (30% methanol, 10% acetic acid in water) and scanned using Cy5 laser/emission filters on an Amersham Typhoon imager (Cytiva). Total protein staining was done using Coomassie R-250 (Thermo Scientific, CAS 6104-58-1).

Adjustment of image brightness and contrast and band intensity quantification was performed with Fiji[17]. Statistical comparison of mean band intensities between uninhibited and inhibited enzymes was done using two-tailed t-tests with three replicates per group.

**Relative Activity of *Mtb* Enzymes**

The relative enzymatic activity was determined by nitrocefin (EMD Millipore, CAS 41906-86-9) hydrolysis. Enzymes from the small-scale purification were concentrated using 10k MWCO centrifugal filters (Amicon, 0.5 mL). A BCA assay was performed to quantify protein concentration. Samples were pre-equilibrated to 37 °C for 15 min in 50 mM Tris-Cl (pH 7.2 at 37 °C), 150 mM NaCl. Each enzyme [10 µM (DacB, Rv1367c, Rv1723, Rv2257c, DapE, Rv0309), 8 µM MurI, 1.4 µM LipD, and 0.1 µM BlaC] was combined with nitrocefin (200 µM) in a flat-bottom, half-well 96-well transparent plate (Corning 3695). Immediately after mixing, nitrocefin hydrolysis was monitored at 486 nm on a microplate reader (Tecan, 1 min intervals, 2 hr duration, 37 °C). Amount of nitrocefin hydrolyzed was determined by converting the absorbance to concentration using Beer’s law (ε = 20 500 M^−1^ cm^−1^; ℓ = 0.32 cm). Initial velocities were taken from the first 0-600 s (or 600-1200 s [Rv0309]) and were divided by the amount of enzyme (in µM) to determine the relative activity of each enzyme. Relative activities were plotted in Graphpad Prism (version 10.2.0) and error was calculated as the standard error of technical triplicates (n=3).

**Kinetics Analysis: Rv1723, Rv2257c, and Rv0309**

The β-lactamase activity of Rv1723, MBP-Rv2257c and Rv0309 (large-scale purifications) was assayed using nitrocefin. Buffer (control) or enzyme was combined with nitrocefin (100-1200 µM) in a flat-bottom, 96-well half-well plate (Corning 3695) in 50 mM Tris-Cl, 150 mM NaCl, pH 8.0 at 4 °C. The reaction (40 µL) was initiated by addition of the enzyme: Rv1723 (35 μM), MBP-Rv2257c (35 μM), or Rv0309 (30 μM). Nitrocefin hydrolysis was monitored spectroscopically (absorbance at 486 nm) on a microplate reader (Tecan; 25-30 sec intervals, 4-10 hr duration, 37 °C). Initial velocities were taken from the first 75-475 s (Rv1723 and Rv2257c) or 750-1500 s (Rv0309) of the reaction and assumed to follow Michaelis-Menten kinetics. Velocities versus substrate concentration were fitted using a non-linear regression in Graphpad Prism (version 10.2.0). Calculations were done using technical triplicates from three (Rv1723 and Rv0309) or four (Rv2257c) independent experiments. A ROUT test (Q=1%) was performed to identify outliers in initial velocities used to calculate the K_M_ for each enzyme, and one was found and omitted from the 400 µM nitrocefin condition for Rv2257c. Error was calculated from a symmetrical (asymptotic) approximate confidence interval to generate a standard error of the mean.

**Inhibition of Enzyme Activity by β-lactams**

The loss of activity of Rv1723 and Rv0309 in the presence of β-lactam inhibitors was assayed using nitrocefin, as described above. Rv1723 and Rv0309 (18 μM) were treated with either buffer only (no treatment) or a 5-fold excess (90 μM) of meropenem, tebipenem, cefoxitin, or amoxicillin (30 min, 37 °C). Samples were then combined with nitrocefin (600 μM [Rv1723] or 1400 μM [Rv0309]) in a flat-bottom, 96-well half-well plate (Corning 3695). Immediately after mixing, nitrocefin hydrolysis was monitored at 486 nm on a Tecan microplate reader (15 min intervals, 3 hr duration, 37 °C). Absorbance values at time 0 were subtracted from the absorbance values at 2 hr, then converted to amount of hydrolyzed nitrocefin using Beer’s law. One-way ANOVA with multiple comparisons tests were performed using Graphpad Prism software to determine statistically significant difference between groups. All calculations were done using technical triplicates from three separate experiments and error was calculated as the standard error of the mean.

**Native Mass Spectrometry of Enzyme-Inhibitor Complexes**

Rv0309 was used without further purification. Rv1723 was further purified by size-exclusion chromatography using a Superdex 200 10/300 GL column (Cytiva). Immediately prior to shipment on ice packs, purified enzymes were buffer exchanged (twice) into 200 mM ammonium acetate, pH 8.0 using MicroBioSpin P-6 columns (BioRad). Antibiotic inhibitors were made fresh at 10 mM and then incubated with each enzyme at a 25-molar excess at 37 °C for 1 hr. To observe inhibitor binding, native MS experiments were performed using the Synapt G2si quadrupole ion mobility time-of-flight mass spectrometer (Waters). Static nanoelectrospray was used to infuse each sample using in-house pulled borosilicate glass capillaries (ID 0.78 mm, item no. BF100-78-10, Sutter Instruments) using a Sutter Instrument pipet puller P-1000 (Novato). Data were collected in positive ion mode, using 4.0 V CID, sampling cone voltage 150 V, source offset 80 V, capillary voltage ∼1.5 kV, and 60 °C source temperature. Native MS data analysis was performed using UniDec deconvolution software[18]. Spectra are provided in **Figure S6**.

**Supplementary Data**

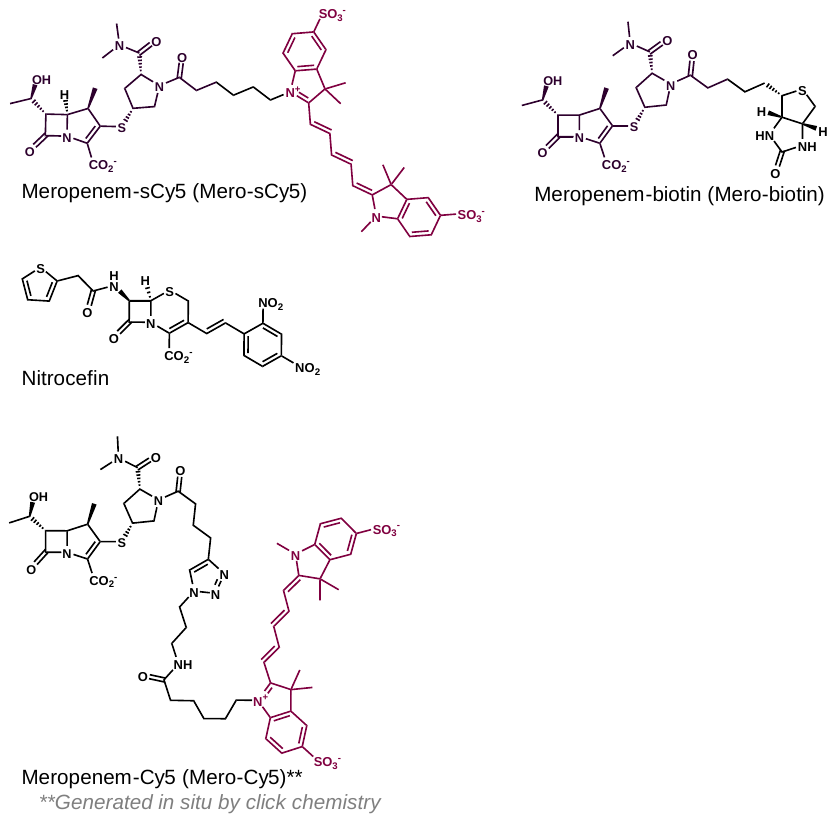

**Scheme S1. Structure of new ABPs (Mero-sCy5 and Mero-biotin), nitrocefin, and previously described Mero-Cy5[****19****].**

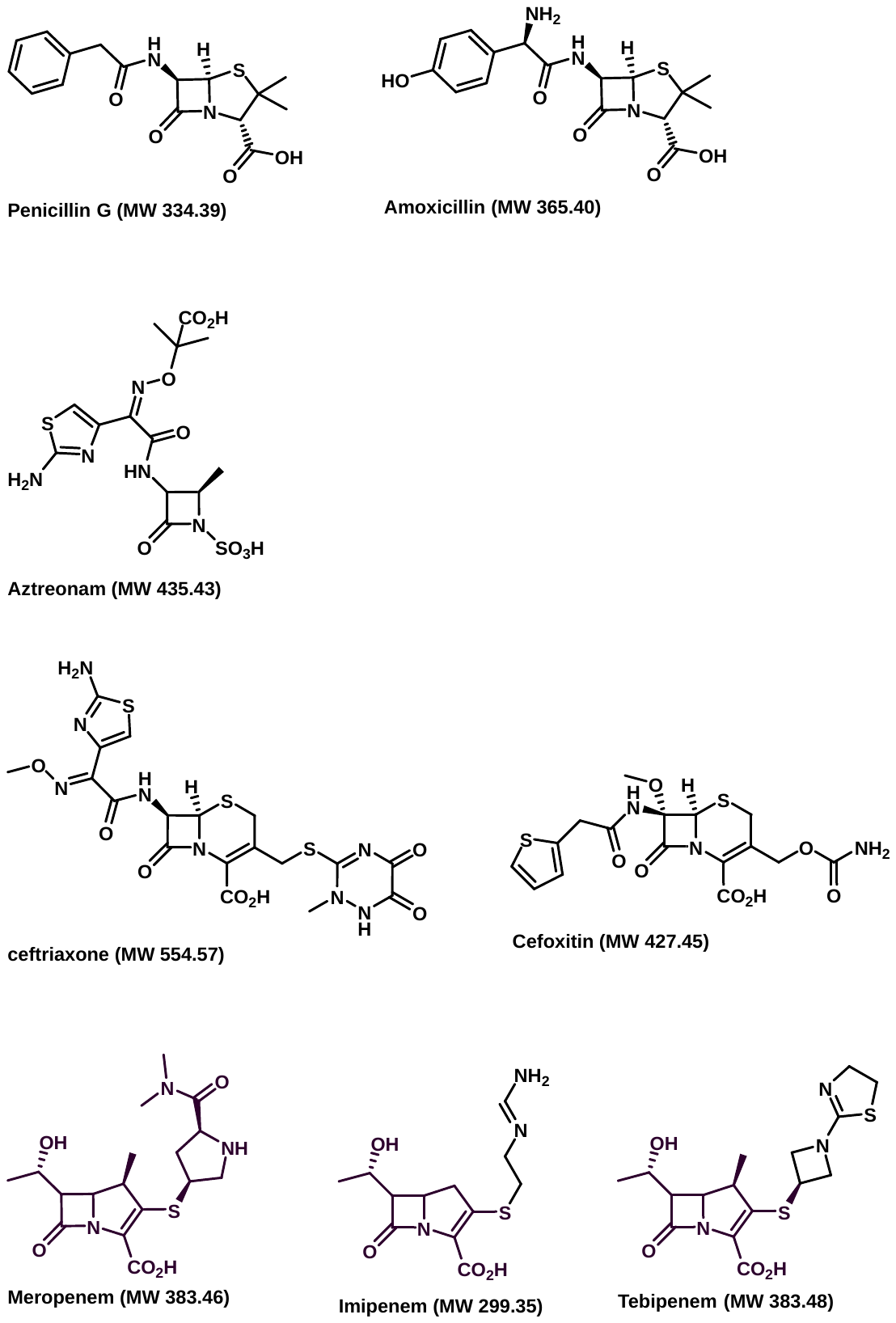

**Scheme S2. Structures of antibiotics used in the current work.**

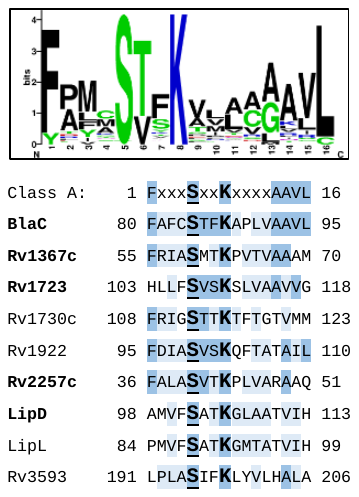

**Figure S1. β-lactamase active site alignment.** Class A β-lactamase motif sequence alignment from Prosite (PS00146; prosite.expasy.org). Conservation of the Class A β-lactamase motif for BlaC, Rv1367c, and uncharacterized enzymes identified by ABPP. Color-coding indicates sequence similarity to the consensus and the active site Ser (**S**) is underlined. Enzymes in bold were characterized in the current work.

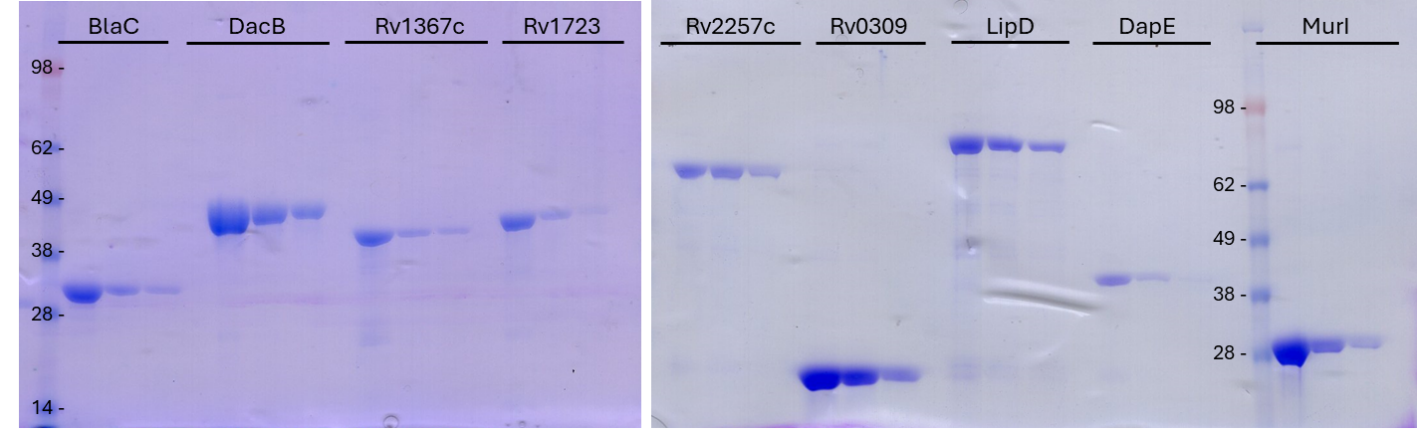

**Figure S2. Small scale purification of enzymes**. Soluble enzymes were purified by immobilized metal affinity chromatography and eluted thrice in 50 mM NaH_2_PO_4_, 300 mM NaCl, 250 mM imidazole, pH 8.0. The first three elution fractions were analyzed for purity by SDS-PAGE. The first elution was used in subsequent experiments. Calculated molecular weight (kDa) of expressed enzymes were as follows: BlaC = 31.41; DacB = 45.81; Rv1367c = 43.41, Rv1723 = 43.77; MBP-Rv2257c = 73.59; Rv0309 = 22.33; MBP-LipD = 94.03, DapE = 39.68; MurI = 30.78.

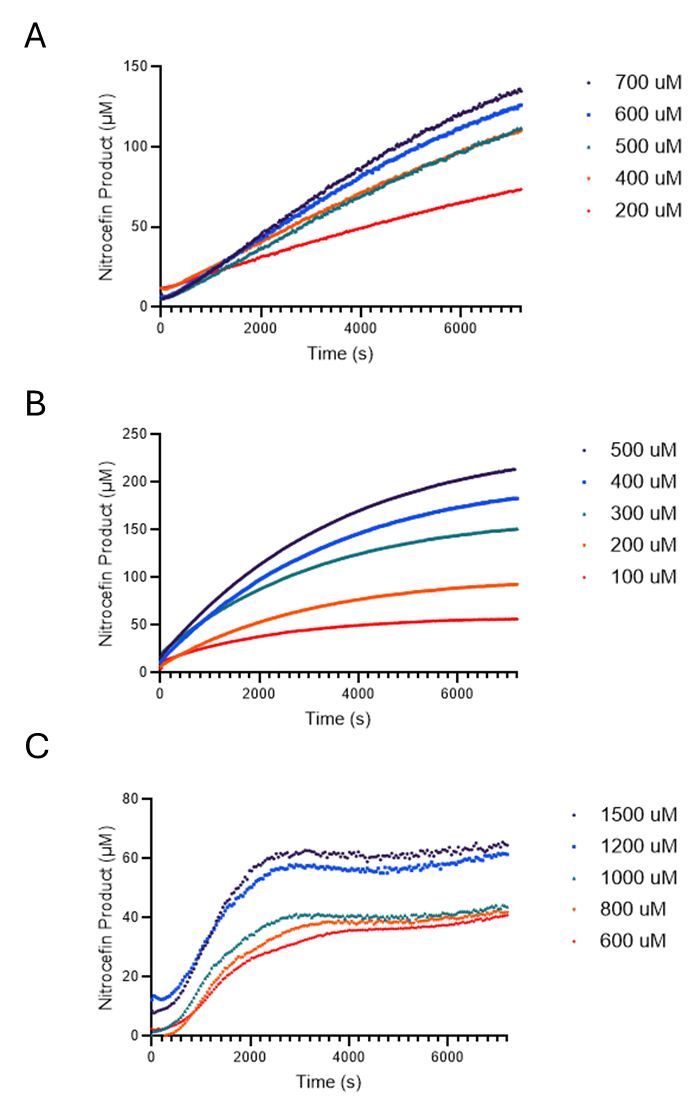

**Figure S3. Hydrolysis of nitrocefin over time by purified enzymes.** Representative traces of a single replicate are shown for Rv1723 (**A**), Rv2257 (**B**), and Rv0309 (**C**). Formation of product was monitored continuously at 37°C in Tris Buffer (pH 7.2).

Figure S4. ^1^H-NMR of meropenem-biotin [mero-biotin] (400 MHz; D_2_O).

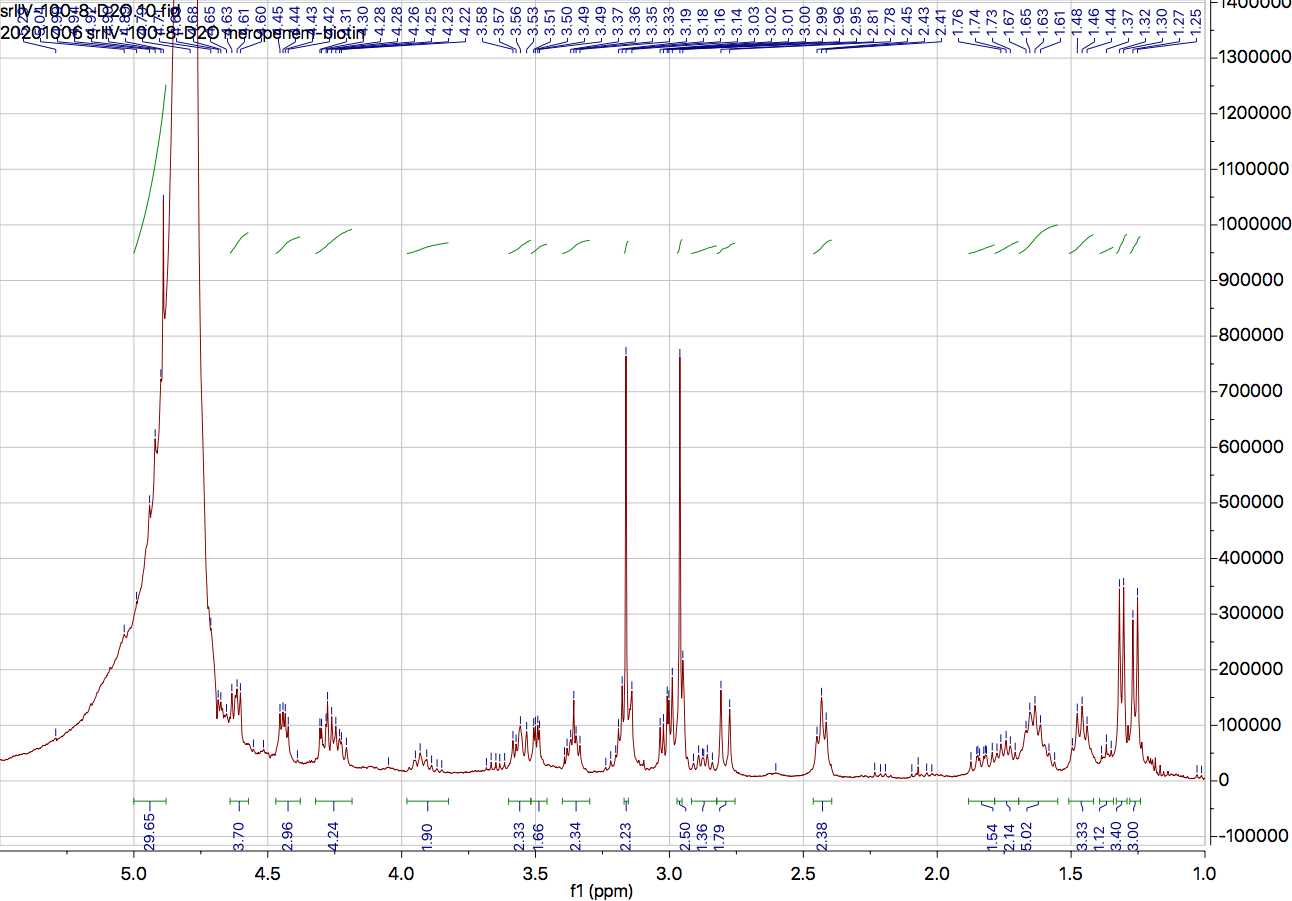

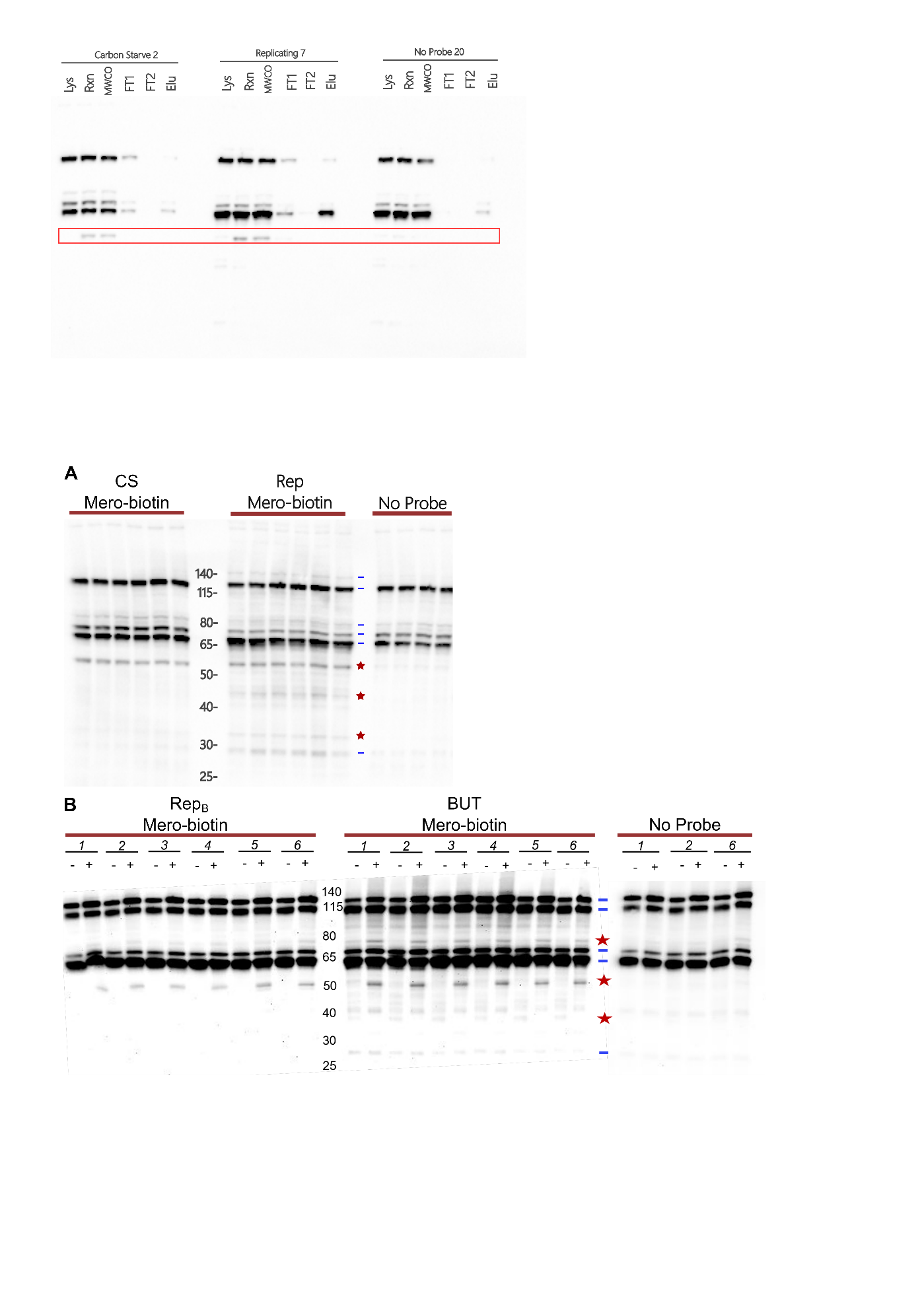

**Figure S5. Western blot analysis of ABPP samples.** Probed (Mero-biotin) and unprobed (No Probe) total protein samples from CS and matched Rep (**A**) and BUT and matched Rep_B_ (**B**) were analyzed by anti-biotin western blot. Samples were normalized by protein amount. Endogenously biotinylated proteins (identified by presence in probed and no-probe samples) are marked with a blue dash. Probe labeled proteins are marked with a red star.

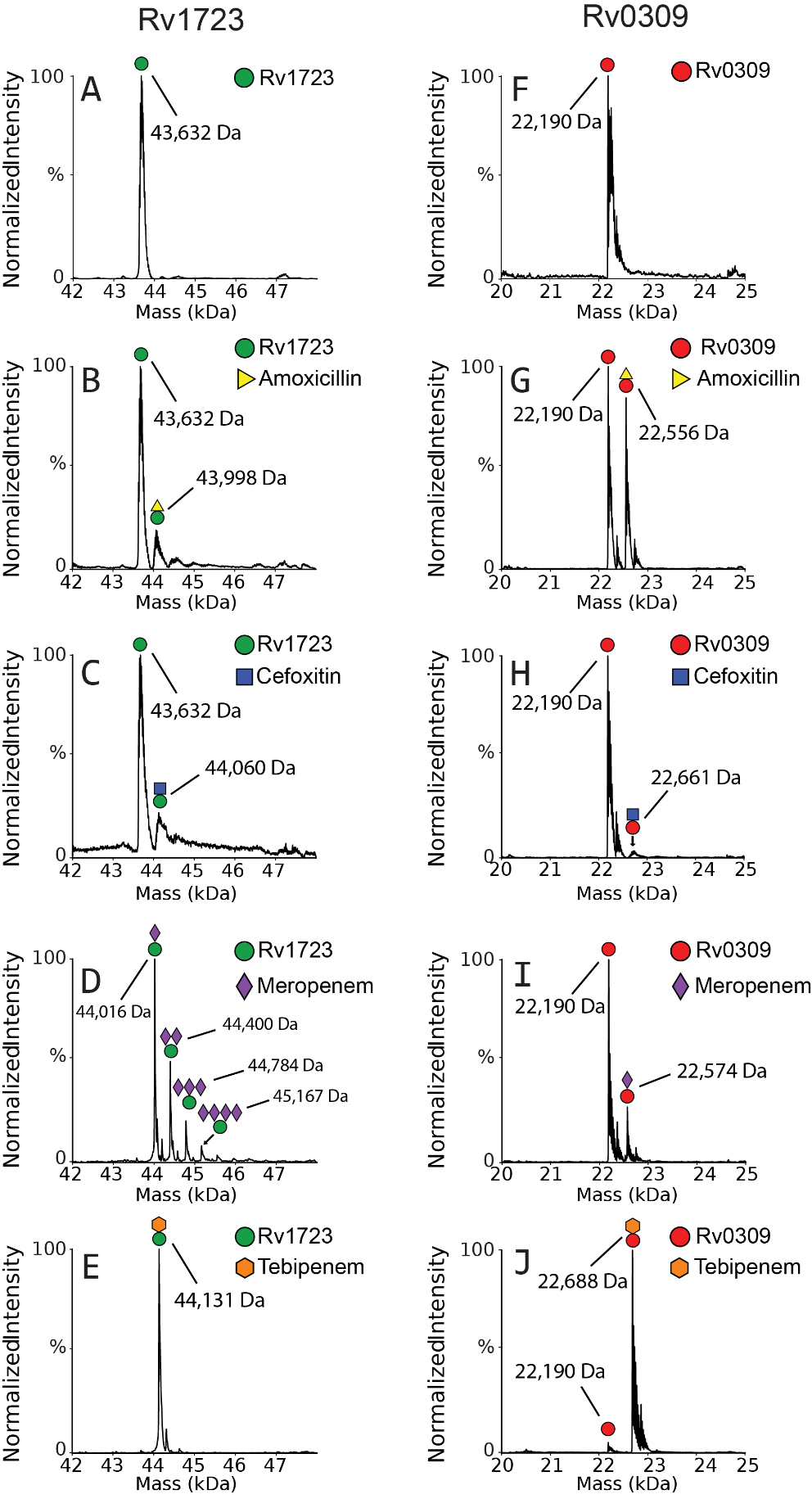

**Figure S6.** **Native MS analysis of Rv1723 and Rv0309 binding of meropenem, tebipenem, amoxicillin, and cefoxitin**. Deconvolved mass spectra for Rv1723 alone and with each inhibitor are shown in **A –** **E**. Deconvolved mass spectra for Rv0309 control and with each inhibitor are shown in **F – J**. Although some binding was observed for Rv1723 with amoxicillin (**B**) and cefoxitin (**C**), significant binding was observed for meropenem (**D**) and tebipenem (**E**). Rv0309 displayed significant binding with amoxicillin (**G**) and tebipenem (**J**), partial binding with meropenem (**I**) and nearly no binding with cefoxitin (**H**).

**Supplementary Tables**

**Table S1: *Mtb* proteins with endogenous biotinylation or biotin binding.**

| **Name** | **Locus ID** | **MW**  **(kDa)** | **Mycobrowser Function** |
| --- | --- | --- | --- |
| AccA1 | Rv2501c | 70.59 | Biotin carboxyl carrier protein and biotin carboxyltransferase. |
| AccA2 | Rv0973c | 70.74 | Biotin carboxyl carrier protein and biotin carboxyltransferase. |
| AccA3 | Rv3285 | 63.75 | Biotin carboxyl carrier protein and biotin carboxyltransferase. |
| AccD1 | Rv2502c | 56.75 | Component of the acetyl coenzyme A carboxylase complex. |
| AccD2 | Rv0974C | 56.22 | Involved in fatty acid metabolism. |
| AccD3 | Rv0904C | 51.74 | Component of the acetyl coenzyme A carboxylase complex. |
| AccD4 | Rv3799c | 56.65 | Key enzyme in the catabolic pathway of odd-chain fatty acids, isoleucine, threonine, methionine, and valine. |
| AccD5 | Rv3280 | 59.35 | Key enzyme in the catabolic pathway of odd-chain fatty acids, isoleucine, threonine, methionine, and valine. |
| AccD6 | Rv2247 | 50.14 | Involved in fatty acid biosynthesis (mycolic acids synthesis). |
| AccE5 | Rv3281 | 19.01 | Involved in long-chain fatty acid synthesis. |
| BioA | Rv1568 | 46.32 | Involved in bioconversion of pimelate into dethiobiotin. Supposedly involved in stationary-phase survival. |
| BioB | Rv1589 | 37.52 | Involved in biotin synthesis. |
| BioD | Rv1570 | 22.44 | Involved in bioconversion of pimelate into dethiobiotin. |
| BioF1 | Rv1569 | 40.03 | Involved in biotin biosynthesis. |
| BioF2 | Rv0032 | 86.24 | Could be involved in biotin biosynthesis. |
| BirA | Rv3279c | 28.09 | Biotin-operon repressor and enzyme that synthesizes the co-repressor, acetyl-CoA:carbon-dioxide ligase. Activates biotin to form biotinyl-5'-adenylate and transfers the biotin moiety to biotin-accepting proteins. |
| BisC | Rv1442 | 83.39 | This enzyme may serve as a scavenger, allowing the cell to utilize biotin sulfoxide as a biotin source. |
| PCA | Rv2967c | 120.39 | Involved in gluconeogenesis and lipogenesis. Catalyzes a 2-step reaction, involving the ATP-dependent carboxylation of the covalently attached biotin in the first step and the transfer of the carboxyl group to pyruvate in the second. |
| Tb7.3 | Rv3221 | 73.06 | Function unknown. (Biotinylated protein TB7.3). |

**Table S2:** *See corresponding Excel file “****ESI Table S2_ABPP hits****”.*

**Table S3:** *See corresponding Excel file “****ESI Table S3_ABPP hit overlap****”*.

**Table S4: *M. tuberculosis* mc^2^6020 Culture Media Recipes.**

| **Medium** | **Component** | **Supplier** | **Final Concentration** |
| --- | --- | --- | --- |
| **7H9/OADC-KPC** | Middlebrook 7H9 broth | BD Difco #271310 | 4.7 g per 1 L |
|  | Middlebrook OADC | BD Difco #212351 | 10% |
|  | Glycerol (molecular biology grade) | Fisher | 0.5% |
|  | Tween 80 | Sigma | 0.05% |
|  | Casamino acids | Gibco #223120 | 0.2% |
|  | Pantothenate | Sigma | 24 μg/mL |
|  | Lysine | Sigma | 80 μg/mL |
|  | H_2_O |  |  |
| **7H9/Tx-KP** | Middlebrook 7H9 broth | BD Difco #271310 | 4.7 g per 1 L |
|  | Tyloxapol | Sigma | 0.05% |
|  | Pantothenate | Sigma | 24 μg/mL |
|  | Lysine | Sigma | 80 μg/mL |
|  | H_2_O |  |  |
| **7H9/Butyrate-KP** | Middlebrook 7H9 broth | BD Difco #271310 | 4.7 g per 1 L |
|  | Fatty-acid free BSA | RPI #A30075 | 0.5 g per 1 L |
|  | NaCl | Fisher | 100 mM |
|  | Tyloxapol | Sigma | 0.05% |
|  | MOPS, pH 7 | Fisher | 100 mM |
|  | Pantothenate | Sigma | 24 μg/mL |
|  | Lysine | Sigma | 80 μg/mL |
|  | Butyrate | Sigma #303410 | 5 mM |
|  | H_2_O |  |  |
| **7H9/OADC/MOPS-KP** | Middlebrook 7H9 broth | BD Difco #271310 | 4.7 g per 1 L |
|  | Middlebrook OADC | BD Difco #212351 | 10% |
|  | Glycerol (molecular biology grade) | Fisher | 0.5% |
|  | Tween 80 | Sigma | 0.05% |
|  | MOPS, pH 7 |  | 100 mM |
|  | Pantothenate | Sigma | 24 μg/mL |
|  | Lysine | Sigma | 80 μg/mL |
|  | H_2_O |  |  |

**Table S5: Summary of *Mtb* “hit” protein constructs.**

| **Enzyme** | **Locus ID** | **UniProt ID** | **MW (kDa)** | **Amino Acid Sequence^‡^** | **Vector name** |
| --- | --- | --- | --- | --- | --- |
| **BlaC** | Rv2068c | [P9WKD3](http://www.uniprot.org/uniprot/P9WKD3) | 31.41 | MGSSHHHHHHSSGENLYFQGHGADLADRFAELERRYDARLGVYVPATGTTAAIEYRADERFAFCSTFKAPLVAAVLHQNPLTHLDKLITYTSDDIRSISPVAQQHVQTGMTIGQLCDAAIRYSDGTAANLLLADLGGPGGGTAAFTGYLRSLGDTVSRLDAEEPELNRDPPGDERDTTTPHAIALVLQQLVLGNALPPDKRALLTDWMARNTTGAKRIRAGFPADWKVIDKTGTGDYGRANDIAVVWSPTGVPYVVAVMSDRAGGGYDAEPREALLAEAATCVAGVLA | pET28a_His6-TEV-BlaC(41-307) |
| **DacB** | Rv3627c | [O06380](http://www.uniprot.org/uniprot/O06380) | 45.81 | MGSSHHHHHHSSGENLYFQGHGGHRAGVRAPAPPPRPPTVKAGVVPVADTAATPSAAGVTAALAVVAADPDLGKLAGRITDALTGQELWQRLDDVPLVPA**STNK**ILTAAAALLTLDRQARISTRVVAGGQNPQGPVVLVGAGDPTLSAAPPGQDTWYHGAARIGDLVEQIRRSGVTPTAVQVDASAFSGPTMAPGWDPADIDNGDIAPIEAAMIDAGRIQPTTVNSRRSRTPALDAGRELAKALGLDPAAVTIASAPAGARQLAVVQSAPLIQRLSQMMNASDNVMAECIGREVAVAINRPQSFSGAVDAVTSRLNTAHIDTAGAALVDSSGLSLDNRLTARTLDATMQAAAGPDQPALRPLLDLLPIAGGSGTLGERFLDAATDQGPAGWLRAKTGSLTAINSLVGVLTDRSGRVLTFAFISNEAGPNGRNAMDALATKLWFCGCTT | pET28a_His6-TEV-DacB(35-461) |
| **Rv1367c** | Rv1367c | [P9WLZ3](http://www.uniprot.org/uniprot/P9WLZ3) | 43.41 | MGSSHHHHHHSSGENLYFQGHMNLDGNQASIREVCDAGLLSGAVTMVWQREKLLQVNEIGYRDIDAGVPMQRDTLFRIA**SMTK**PVTVAAAMSLVDEGKLALRDPITRWAPELCKVAVLDDAAGPLDRTHPARRAILIEDLLTHTSGLAYGFSVSGPISRAYQRLPFGQGPDVWLAALATLPLVHQPGDRVTYSHAIDVLGVIVSRIEDAPLYQIIDERVLGPAGMTDTGFYVSADAQRRAATMYRLDEQDRLRHDVMGPPHVTPPSFCNAGGGLWSTADDYLRFVRMLLGDGTVDGVRVLSPESVRLMRTDRLTDEQKRHSFLGAPFWVGRGFGLNLSVVTDPAKSRPLFGPGGLGTFSWPGAYGTWWQADPSADLILLYLIQHCPDLSVDAAAAVAG | pET28a_His6-TEV-Rv1367c(1-377) |
| **Rv1723** | Rv1723 | [P71981](http://www.uniprot.org/uniprot/P71981) | 43.77 | MGSSHHHHHHSSGENLYFQGHMSGGVPAGLALDNWLSSPYSHWAFQHVEDFMPTTVIARGTEPVVTLPADNAPIADIGLTSTDGIATTVGAVMAATATDGWAVAHRGALVAEQYLDGLGPRTRHLLF**SVSK**SLVAAVVGALHGAGAIELDAPVTAYVPALADCGYAGATVRHLLDMRSGVAFSENYDDPAAEIHVREQVIGWAPKRGPDLPATLRDYLLTLRRKSAHGGPFEYRSCETDVLGWICEAAAGQPMPELMSELLWSRIGAQCDATIALDVAGAAGTGIFDGGISACLTDMIRFGSLYLRDGVSLAGQQVVPAAWIADTFDGGPDSRQAFAASPDDNPMPGGMYRNQVWFPYPGSNVALCVGMCGQLIYVNRAAEVVAAKLSTQPHSHEPHMLDTLRAFDAVAHELSG | pET28a_His6-TEV-Rv1723(1-393) |
| **MBP-Rv2257c** | Rv2257c | [O53531](http://www.uniprot.org/uniprot/O53531) | 73.59 | MHHHHHHHGPGGKIEEGKLVIWINGDKGYNGLAEVGKKFEKDTGIKVTVEHPDKLEEKFPQVAATGDGPDIIFWAHDRFGGYAQSGLLAEITPDKAFQDKLYPFTWDAVRYNGKLIAYPIAVEALSLIYNKDLLPNPPKTWEEIPALDKELKAKGKSALMFNLQEPYFTWPLIAADGGYAFKYENGKYDIKDVGVDNAGAKAGLTFLVDLIKNKHMNADTDYSIAEAAFNKGETAMTINGPWAWSNIDTSKVNYGVTVLPTFKGQPSKPFVGVLSAGINAASPNKELAKEFLENYLLTDEGLEAVNKDKPLGAVALKSYEEELAKDPRIAATMENAQKGEIMPNIPQMSAFWYAVRTAVINAASGRQTVDEALKDAQTNSSSNNNNNNNNNNLGIDTTENLYFQGAMDPEFMTALEVLGGWPVPAAAAAVIGPAGVLATHGDTARVFALA**SVTK**PLVARAAQVAVEEGVVNLDTPAGPPGSTVRHLLAHTSGLAMHSDQALARPGTRRMYSNYGFTVLAESVQRESGIEFGRYLTEAVCEPLGMVTTRLDGGPAAAGFGATSTVADLAVFAGDLLRPSTVSAQMHADATTVQFPGLDGVLPGYGVQRPNDWGLGFEIRNSKSPHWTGECNSTRTFGHFGQSGGFIWVDPKADLALVVLTARDFGDWALDLWPAISDAVLAEYT | pParallel_His7-MBP-TEV-Rv2257c(1-272) |
| **DapE** | Rv1202 | [P9WHS9](http://www.uniprot.org/uniprot/P9WHS9) | 39.68 | MGSSHHHHHHSSGENLYFQGHMLDLRGDPIELTAALIDIPSESRKEARIADEVEAALRAQASGFEIIRNGNAVLARTKLNRSSRVLLAGHLDTVPVAGNLPSRRENDQLHGCGAADMKSGDAVFLHLAATLAEPTHDLTLVFYDCEEIDSAANGLGRIQRELPDWLSADVAILGEPTAGCIEAGCQGTLRVVLSVTGTRAHSARSWLGDNAIHKLGAVLDRLAVYRARSVDIDGCTYREGLSAVRVAGGVAGNVIPDAASVTINYRFAPDRSVAAALQHVHDVFDGLDVQIEQTDAAAGALPGLSEPAAKALVEAAGGQVRAKYGWTDVSRFAALGIPAVNYGPGDPNLAHCRDERVPVGNITAAVDLLRRYLGG | pET28a_His6-TEV-DapE(1-354) |
| **Rv0309** | Rv0309 | [O07236](http://www.uniprot.org/uniprot/O07236) | 22.33 | MGSSHHHHHHSSGENLYFQGHMSNPWFANSVGNATQVVSVVGTGGSTAKMDVYQRTAAGWQPLKTGITTHIGSAGMAPEAKSGYPATPMGVYSLDSAFGTAPNPGGGLPYTQVGPNHWWSGDDNSPTFNSMQVCQKSQCPFSTADSENLQIPQYKHSVVMGVNKAKVPGKGSAFFFHTTDGGPTAG**C**VAIDDATLVQIIRWLRPGAVIAIAK | pET28a_His6-TEV-Rv0309(30-218) |
| **MurI** | Rv1338 | [P9WPW9](http://www.uniprot.org/uniprot/P9WPW9) | 30.78 | MGHHHHHHENLYEQ*SHMNSPLAPVGVFDSGVGGLTVARAIIDOLPDEDIVYVGDTGNGPYGPLTIPEIRAHALAIGDDLVGRGVKALVIACNSASSACLRDARERYQVPVVEVILPAVRRAVAATRNGRIGVIGTRATITSHAYQDAFAAARDTEITAVACPRFVDFVERGVTSGRQVLGLAQGYLEPLQRAEVDTLVLGCTHYPLLSGLIQLAMGENVTLVSSARETAKEVVRVLTEIDLLRPHDAPPATRIFEATGDPBAFTKLAARFLGPVLGGVQPVHPSRIH | pET28a_His6-TEV-MurI(1-271) |
| **MBP-LipD** | Rv1923 | [P95290](http://www.uniprot.org/uniprot/P95290) | 94.03 | MHHHHHHHGPGGKIEEGKLVIWINGDKGYNGLAEVGKKFEKDTGIKVTVEHPDKLEEKFPQVAATGDGPDIIFWAHDRFGGYAQSGLLAEITPDKAFQDKLYPFTWDAVRYNGKLIAYPIAVEALSLIYNKDLLPNPPKTWEEIPALDKELKAKGKSALMFNLQEPYFTWPLIAADGGYAFKYENGKYDIKDVGVDNAGAKAGLTFLVDLIKNKHMNADTDYSIAEAAFNKGETAMTINGPWAWSNIDTSKVNYGVTVLPTFKGQPSKPFVGVLSAGINAASPNKELAKEFLENYLLTDEGLEAVNKDKPLGAVALKSYEEELAKDPRIAATMENAQKGEIMPNIPQMSAFWYAVRTAVINAASGRQTVDEALKDAQTNSSSNNNNNNNNNNLGIDTTENLYFQGAMDPEFDVAGLPRLAAGTQAAIIHGMAQPPSLLTTDNGLPFGVQGACDSRFTGVIRAFAGLYPGRKFGGGALSVYIDGRQVVDVWTGWSDRQGKVPWTADTGAMVF**SATK**GLAATVIHRLVDRGLLSYDAPVAEYWPEFGANGKSEVTVSDVLRHRSGLAHLKGVDKDEVMDHLLMEQKLAAAPLDRQHGKLAYHAVTYGWLLSGLARAVTGKGMRELFREELARPLNTDGIHLGRPPADSPTKAAQTLLPQAKVPTPLLDFIAPKVAGLSFSGLLGAVYFPGILSLLQDDMPFLDGEVPAVNGVVTARALAKTYGALANDGVIDGTRLLSSQAVRGLTGKSELWPDLNLGLPFTYHQGYQSSPVPGLLEGYGHIGLGGTIGWADPETGSAFGYVHNRLLTLLLFDIGSFAGLAALLNSAVVAARRDDPLEVPHFGAPYSEPRHEQAASGALEACGTKLGTGRRFTTS | pParallel_His7-MBP-TEV-LipD(1-446) |
| **Rv1723 (S107A)** |  |  | 43.76 | MGSSHHHHHHSSGENLYFQGHMSGGVPAGLALDNWLSSPYSHWAFQHVEDFMPTTVIARGTEPVVTLPADNAPIADIGLTSTDGIATTVGAVMAATATDGWAVAHRGALVAEQYLDGLGPRTRHLLF**AVSK**SLVAAVVGALHGAGAIELDAPVTAYVPALADCGYAGATVRHLLDMRSGVAFSENYDDPAAEIHVREQVIGWAPKRGPDLPATLRDYLLTLRRKSAHGGPFEYRSCETDVLGWICEAAAGQPMPELMSELLWSRIGAQCDATIALDVAGAAGTGIFDGGISACLTDMIRFGSLYLRDGVSLAGQQVVPAAWIADTFDGGPDSRQAFAASPDDNPMPGGMYRNQVWFPYPGSNVALCVGMCGQLIYVNRAAEVVAAKLSTQPHSHEPHMLDTLRAFDAVAHELSG | pET28a_His6-TEV-Rv1723(S107A) |
| **Rv0309 (C193G)** |  |  | 22.28 | MGSSHHHHHHSSGENLYFQGHMSNPWFANSVGNATQVVSVVGTGGSTAKMDVYQRTAAGWQPLKTGITTHIGSAGMAPEAKSGYPATPMGVYSLDSAFGTAPNPGGGLPYTQVGPNHWWSGDDNSPTFNSMQVCQKSQCPFSTADSENLQIPQYKHSVVMGVNKAKVPGKGSAFFFHTTDGGPTAG**G**VAIDDATLVQIIRWLRPGAVIAIAK | pET28a_His6-TEV-Rv0309(C193G) |

**‡	Key:** His tag, TEV recognition site, maltose-binding protein

Class A beta-lactamase motif: **SxxK**

LDT-catalytic domain cysteine: **C**

**Table S6: Primers for site-directed mutagenesis.**

| **Name** | **Sequence** (5′ → 3′, mutation site) | **Type** | **Description** |
| --- | --- | --- | --- |
| Rv0309_C193G_FWD_1 | GTCCGACTGCGGGtgGCGTTGCGATC | SDM | Forward primer for substitution of Rv0309 C^193^ to G. |
| Rv0309_C193G_REV_1 | CGCcaCCCGCAGTCGGACCGCCGTCAG | SDM | Reverse primer for substitution of Rv0309 C^193^ to G. |
| Rv1723_S107A_FWD | CCTATTGTTTgcgGTCTCCAA ATCC | SDM | Forward primer for substitution of Rv1723 S^107^ to A. |
| Rv1723_S107A_REV | TGACGGGTACGTGGG | SDM | Reverse primer for substitution of Rv1723 S^107^ to A. |
